# Stalling of elongating Pol II triggers Ser7 phosphorylation *in trans* to drive transcription recovery

**DOI:** 10.64898/2025.12.04.692291

**Authors:** Iván Shlamovitz, Yu Bao, Tea Toteva, Alastair Crisp, Shutong Ye, Steven W. Wingett, Roberta Cacioppo, Anna Edmondson, Daniela Castiblanco, Michael A. Boemo, Ana Tufegdžić Vidaković

## Abstract

DNA is scattered with obstacles that stall RNA polymerase II (Pol II) and block the production of full-length transcripts. Two mechanisms are known to resolve stalled Pol II: transcription-coupled nucleotide excision repair (TC-NER) and the “last resort” Pol II ubiquitylation–degradation pathway. Here, we uncover a third, distinct mechanism that alerts incoming Pol II molecules to roadblocks ahead and primes them for efficient elongation. We show that transcription stalling triggers GSK3-mediated phosphorylation of Ser7 residues (Ser7^P^) on the Pol II C-terminal domain. Unexpectedly, this phosphorylation occurs *in trans*: obstacles in gene bodies induce Ser7^P^ on Pol II complexes at gene beginnings. This modification enables processive transcription while restraining excessive Pol II degradation by the “last resort” pathway. Our findings reveal an adaptive system that preserves transcriptional homeostasis by coordinating polymerase behavior across the gene in response to elongation stress.

## Introduction

The largest subunit of Pol II, RPB1, contains a large, disordered and repetitive C-terminal domain (CTD), composed in humans of 52 repeats of the seven amino-acid motif Tyr1-Ser2-Pro3-Thr4Ser5-Pro6-Ser7. The phosphorylation at four CTD residues – Tyr1^P^, Ser2^P^, Thr4^P^ and Ser5^P^ – regulates key steps of transcription such as release from initiation, promoter-proximal pausing, cotranscriptional RNA processing and transcription termination^1–6^. Furthermore, kinases that target these residues have been identified^7–10^. Remarkably, Ser7 is also prominently phosphorylated *in vivo*^11^, but studies of Ser7^P^ on protein-coding genes are fundamentally lacking and the kinase directed towards this residue remains unclear^12–14^.

As Pol II travels across the gene, it encounters obstacles such as DNA damage, DNA-bound proteins and secondary DNA structures, which impede Pol II progression and cause transcription stalling^15–17^. Just a single stalled Pol II molecule blocks the path of all subsequent polymerases travelling in its wake, thereby completely impairing the expression of the underlying gene^15^. Two pathways are known to resolve stalled Pol II: transcription-coupled nucleotide excision repair (TC-NER)^18–20^, and the so-called “last resort” pathway which targets elongation-stalled Pol II for ubiquitylation and proteasomal degradation^21–24^. One single residue on the Pol II catalytic subunit, RPB1, is a focal point for coordinating these mechanisms: lysine 1268 (K1268)^25,26^. This residue is targeted for ubiquitylation by both TC-NER machinery and the “last resort” pathway, which likely place different types of ubiquitin chains on it, leading to different outcomes^25^. TC-NER initiates when Cockayne Syndrome B translocase (CSB) collides with stalled Pol II in front of it, followed by recruitment of Cockayne Syndrome A (CSA), a substrate-recognition subunit of the E3 ubiquitin ligase CRL4^CSA^, that modifies RPB1 K1268, facilitating the assembly and stabilization of the complex that will carry out DNA repair^27–29^. Alternatively, RPB1 K1268 is targeted by a “last resort” E3 ubiquitin ligase that leads to its destruction, removing stalled Pol II from the site of DNA damage^25^. Multiple E3 ubiquitin ligases have been implicated in the “last resort”, including NEDD4, CRL5^ELOA^ and CRL2^VHL^ – in cells, these enzymes are functionally redundant, as depletion of neither fully abolishes the “last resort”^30–34^. It is also likely that additional “last resort” E3s are yet to be identified^34^.

Both TC-NER and the “last resort” are essential for cells to survive exposure to genotoxins^25,26,35–37^. In humans, mutations in *CSB* and *CSA* cause a severe progeroid disorder Cockayne Syndrome, while the physiological consequences of losing the “last resort” pathway are unclear due to the redundant and elusive nature of the putative E3s involved^34,36,37^.

Apart from triggering TC-NER and the “last resort”, transcription stalling also causes Pol II hyper-phosphorylation, but the function of this phenomenon has remained unknown^38^. In this study, we reveal that this hyper-phosphorylation occurs specifically at Pol II CTD Ser7. We find that Ser7 hyper-phosphorylation is a universal hallmark of stalled transcription, occurring in response to any type of obstacle that blocks transcription elongation. Remarkably, Ser7 is not hyper-phosphorylated *in cis* – directly on stalled Pol II, but rather *in trans*, at the remote part of the gene, on polymerases entering the early elongation zone. Ser7^P^ constitutes a separate mechanism from TC-NER and the “last resort”, and all these three pathways are required for cells to recover from transcription stalling. Functionally, Ser7^P^ inhibits excessive activity of the “last resort”, salvaging a minimal reserve of Pol II molecules needed for transcription recovery after genotoxic exposure. Using biochemistry, genomics and mathematical modelling, we uncover mechanistic basis of the Ser7^P^ pathway: it is triggered by Pol II backtracking, and the kinase responsible is GSK3. Overall, our results reveal an adaptive system whereby polymerases “communicate” between remote parts of the gene, to maintain transcriptional homeostasis in the face of roadblocks on genes.

## Results

### Ser7^P^ is required for productive Pol II elongation

Ser7^P^ has been implicated in regulating the expression of small nuclear (sn)RNAs^14^, yet snRNAs represent less than 0.5% of all Pol II-transcribed (Class II) genes in a typical human cell type, such as HEK293, a line used throughout this study (Figure S1A). To understand the distribution of Ser7^P^ across the genome we mapped the occupancy of total and Ser7^P^ Pol II using double-crosslinking ChIP-seq (dxChIP-seq). Only a minor fraction of the Ser7^P^ (0.6%) and total Pol II (0.5%) signals correspond to snRNA genes (Figure 1A), showing that most Ser7^P^ Pol II molecules in a cell are located on protein-coding and other non-coding RNA genes, where the function of Ser7^P^ is unknown.

**Figure 1.**
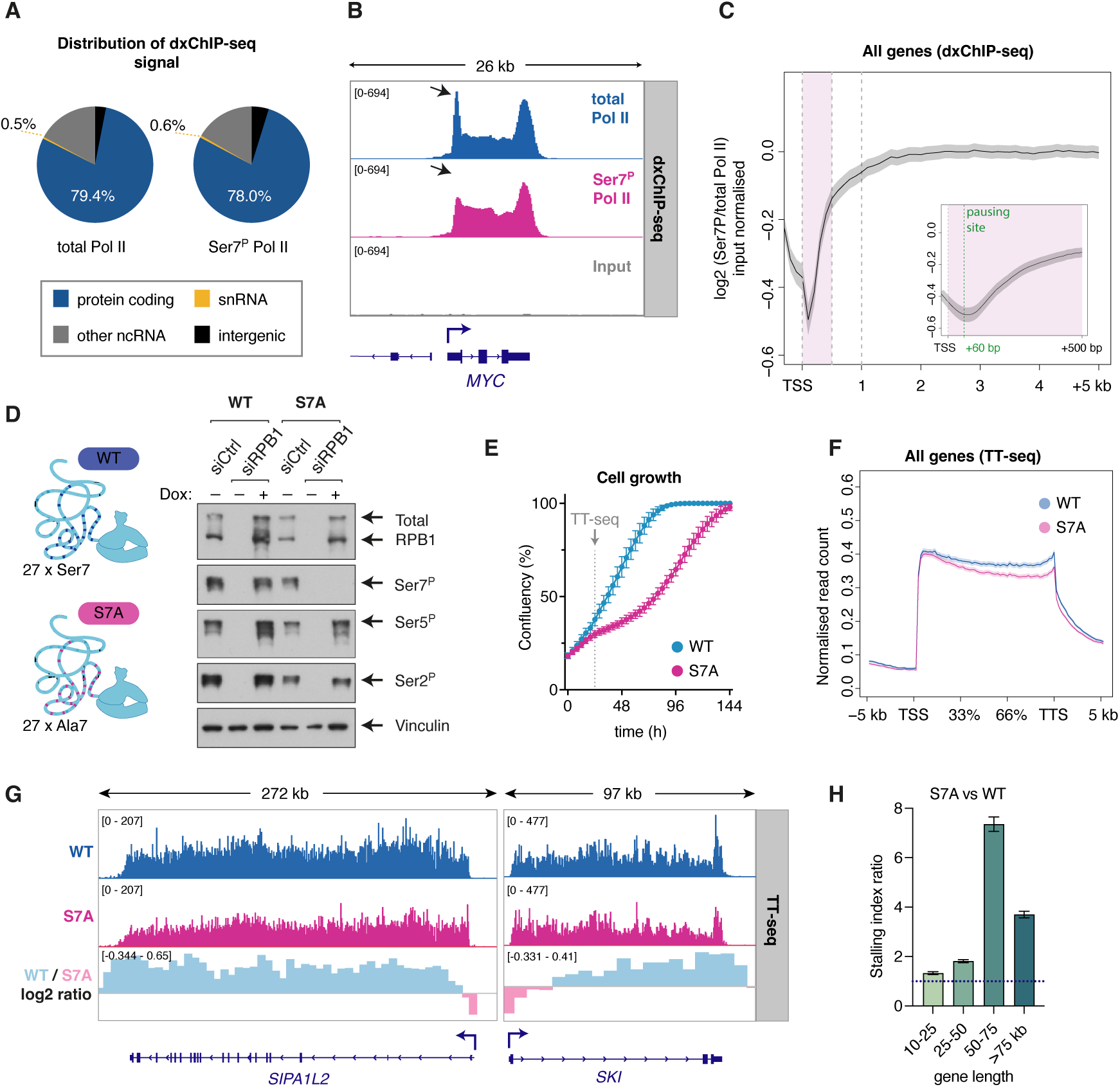
Pol II CTD Ser7 phosphorylation is required for productive elongation. **(A)** Pie charts showing the distribution of total Pol II and Ser7^P^ Pol II signals, detected and quantified by dxChIP-seq, across different gene categories, in WT cells. **(B)** Example gene (*MYC*), displaying dxChIP-seq data for total Pol II and Ser7^P^ in WT cells. Arrow indicates promoter-proximal paused Pol II peak. **(C)** Metagene plots of dxChIP-seq data in WT cells, showing the log2 ratio of input normalized Ser7^P^ versus total Pol II signals, in the first 5 kb and first 500 bp (inset) of genes. **(D)** Switchover system (left) and its efficiency assessed with Western blot, detecting total RPB1 and its different phospho-forms. WT and S7A switchover cells, transfected with non-targeting siRNA (siCtrl) or siRNA against *POLR2A* (siRPB1), with or without induction of transgenic RPB1 with Doxycycline (Dox). **(E)** Growth of WT and S7A switchover cells, monitored by Incucyte. A representative of the biological triplicate experiment is shown, each with 6 technical replicate wells. Data shown as mean +/– standard error of imaging. Arrow indicates the timepoint corresponding to TT_chem_-seq experiment (in f, g, h). **(F)** Metagene plot of TT_chem_-seq data showing nascent RNA production on all genes, in WT and S7A switchover cells. Data are represented on a relative scale (TSS to TTS). Experiment was performed at a time point indicated with an arrow in (e), when cell densities of WT and S7A were comparable. **(G)** Examples showing nascent RNA (TT_chem_-seq data) on *SIPA1L2* and *SKI* genes, in WT and S7A switchover cells. Lower panel shows log2 ratio of WT to S7A signals, with positive values (light blue) indicating higher signal in WT, and negative values (light pink) indicating higher signal in S7A. **(H)** Comparison of stalling indices derived from TT_chem_-seq data, in S7A and WT cells, for different gene length categories. Values above 1 indicate more Pol II stalling in S7A compared to WT cells (indicated with blue dashed line). All values are statistically significant (p<0.0001, two-tailed t-test).

Both globally and at individual genes, Ser7^P^ is associated with elongating Pol II (Figures 1B and S1B), reminiscent of Ser7^P^ Pol II distribution previously characterized in yeast^39–41^. We next inspected the ratio of Ser7^P^ versus total Pol II signals, which indicated that Ser7^P^ is most actively deposited after Pol II leaves the pausing site and enters early elongation (Figure 1C). A similar conclusion has recently been made based on a newly developed technique, PRO-IP-seq, which mapped occupancy of different Pol II phospho-forms with single nucleotide resolution^42^. These findings suggest that Ser7^P^ has a function in regulating Pol II during the early elongation phase, where Pol II assembles elongation factors and gains speed. If this is the case, we would expect elongation to be impaired when Ser7^P^ is abolished.

To determine the function of Ser7^P^, we abolished it using the switchover strategy (SWO) we previously developed^25^. Briefly, endogenous RPB1 is silenced using highly efficient siRNAs, while transgenic, siRNA-resistant and doxycycline-inducible RPB1 is expressed from a singleintegrated FRT locus at a near endogenous level (Figure S1C). We generated S7A SWO cells harboring Ser7 to Ala7 (S7A) mutations in all 27 CTD repeats that contain serine at position 7, completely impairing Ser7^P^ (Figure 1D). WT SWO cells are used as controls (Figure 1D). S7A SWO cells are viable but have impaired growth (Figure 1E), indicating that Ser7^P^ is required for normal cellular function.

We next investigated nascent RNA production genome-wide by TT_chem_-seq^43^, 24 h after switchover, when WT and S7A cells still have similar densities (Figure 1E). This revealed that Pol II elongation is impaired in S7A cells, both globally and at individual gene level (Figures 1F, 1G and S1D). Compared to the WT, S7A mutant cells display a relatively similar (or even increased) nascent RNA signal at gene beginnings, but progressively declining signal throughout the gene body (Figures 1G and S1D). This effect correlates with the length of the gene, with elongation on longer genes being more dramatically affected by the loss of Ser7^P^ (Figures 1H and S1E). Usage of premature transcription termination sites was also increased in S7A mutants, with the effect more pronounced on longer genes, as evidenced by 3’ mRNA-seq (Figure S1F). Together, these data suggest that Ser7^P^ is required for Pol II to elongate processively and avoid premature termination, particularly on long genes.

### Pol II Ser7 hyper-phosphorylation is a universal hallmark of elongation-stalled transcription

During elongation, a major challenge for Pol II is to overcome obstacles that stall its progression^15^. Endogenous obstacles to Pol II are always present in the genome, and exposure to exogenous genotoxins such as UV irradiation, which causes bulky DNA lesions, increases the burden of roadblocks that Pol II faces during elongation^15^. To test if the effects of Ser7^P^ observed above are related to stalling of elongating Pol II, we induced transcription stalling using UV light. Studying stalled Pol II, however, is challenging due to the activity of the “last resort” pathway that leads to its degradation. To stabilize Pol II in the stalled state, we used RPB1 K1268R mutant, which has been shown to abolish the “last resort” Pol II ubiquitylation-degradation and only moderately affect TC-NER efficiency^25,26^.

When K1268R cells are irradiated with UV, hyper-phosphorylation of RPB1 can be observed (Figure 2A). Importantly, using specific antibodies against CTD modifications revealed that this hyper-phosphorylation occurs specifically at Ser7 residues, while phosphorylation at other CTD residues does not change substantially (Figure 2A). The same conclusion was reached in WT cells when degradation of stalled Pol II was blocked using proteasome inhibitor MG-132 (Figure S2A).

**Figure 2.**
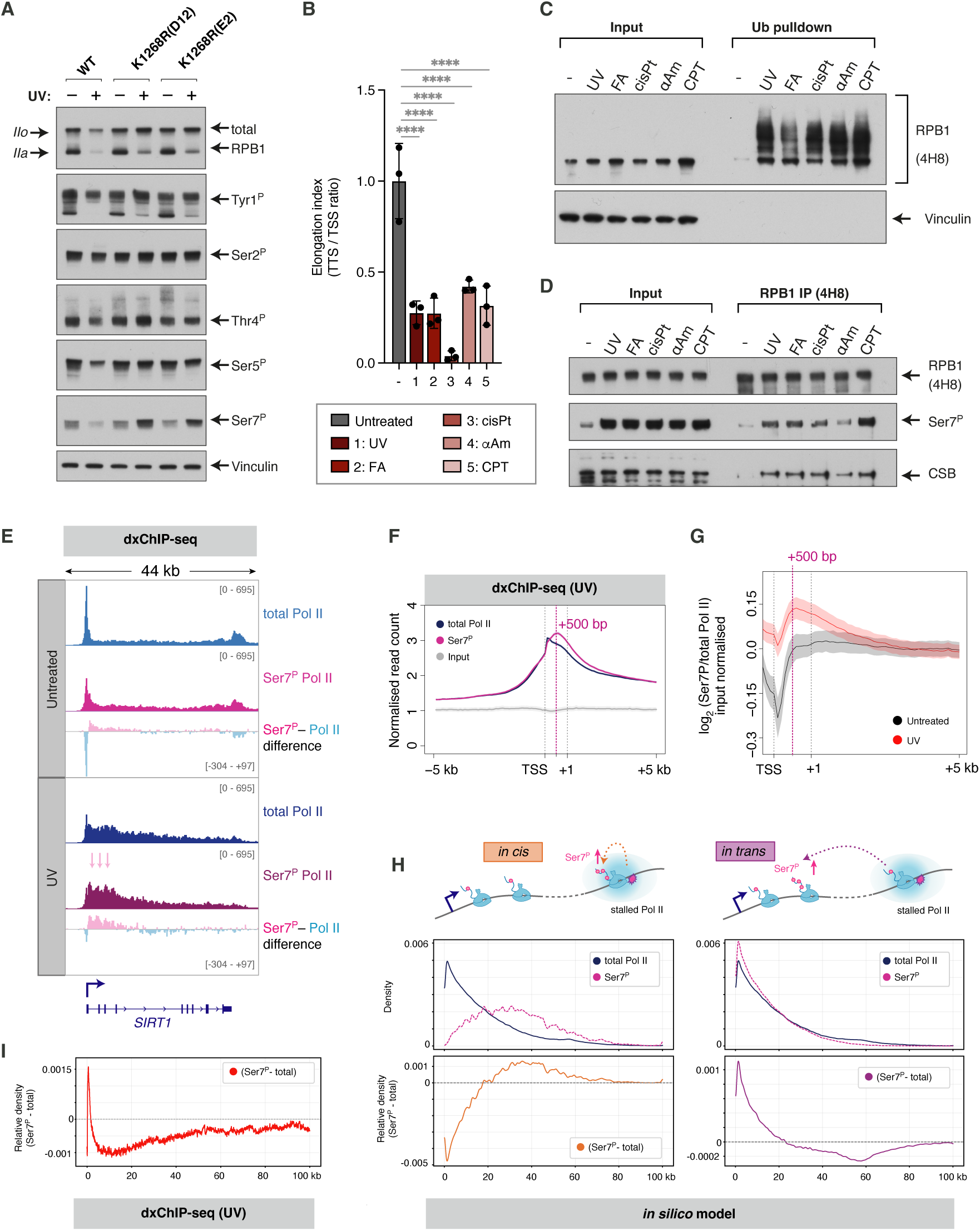
Ser7 hyper-phosphorylation is a universal hallmark of elongation-stalled transcription and it occurs *in trans*. **(A)** Western blot detecting total RPB1 and its different phospho-forms, in WT and two clones of RPB1 K1268R cells (D12 and E2), before and 3 hours post-UV (20 J/m^2^). **(B)** RT-qPCR assay quantifying the Pol II elongation index upon treatment with a panel of stalling agents. Elongation index derived as a ratio of nascent RNA at the end and the beginning of a long, highly expressed gene *EXT1* (see Figure S 2b). Quantification was based on biological triplicates and data represented as mean +/– SD, significance determined using Dunnett’s multiple comparison test, **** indicates p<0.0001. **(C)** Ubiquitin pulldown assay followed by western blot detecting RPB1, before and after treatment with stalling agents, WT cells. **(D)** Immunoprecipitation of RPB1 (4H8 antibody) followed by western blot detecting RPB1, Ser7^P^ and CSB, before and after treatment with stalling agents. WT cells, pre-treated with 5µM MG132 for 3 hours. **(E)** Example of dxChIP-seq data on *SIRT1* gene, for total Pol II, Ser7^P^ and the difference between Ser7^P^ and total Pol II signals, in K1268R cells, before and 3h after UV (20 J/m^2^). Arrows indicate the area of most active Ser7^P^ deposition. **(F)** Metagene plot of dxChIP-seq data, showing the distribution of total and Ser7^P^ Pol II within the first 5 kb from the TSS, 3h after UV (20 J/m^2^), K1268R cells. **(G)** Metagene plot of dxChIP-seq data showing the log2 ratio of Ser7^P^ versus total Pol II signals, within the first 5 kb from the TSS. dxChIP-seq data, K1268R cells, before and 3h after UV (20 J/m^2^). **(H)** Mathematical modelling of *in cis* and *in trans* scenarios for Ser7^P^ deposition. Sketches on top represent assumptions of the model. Simulated total Pol II and Ser7^P^ distributions (middle) and the difference between Ser7^P^ versus total Pol II (bottom) are shown for *in cis* (left) and *in trans* (right) scenarios. The Beacon Calculus results are averaged over 5000 simulations. **(I)** Metagene plot of dxChIP-seq data showing the difference between Ser7^P^ and total Pol II signals, within the first 100 kb from the TSS (on protein coding genes longer than 100 kb). dxChIP-seq data, K1268R cells, 3h after UV (20 J/m^2^).

We next used a panel of agents that are predicted to impose different types of obstacles to elongating Pol II (Figure S2B): UV, which causes intra-strand cyclobutane pyrimidine dimers^17^; formaldehyde (FA), which generates DNA inter-strand and DNA–protein crosslinks^44–47^; cisplatin, which causes DNA intra- and inter-strand crosslinks^48^; alpha-amanitin, which blocks the catalytic center of Pol II inhibiting RNA synthesis^49,50^; and camptothecin (CPT), which covalently traps topoisomerase I on DNA resulting in protein-DNA crosslinks as well as an increase in DNA supercoiling^51^. We validated that these agents indeed stall elongating Pol II using RT-qPCR (Figures 2B and S2C). All types of obstacles triggered the two pathways known to resolve stalled Pol II: the “last resort” poly-ubiquitylation and degradation of RPB1 (Figures 2C and S2D), and recruitment of CSB to Pol II, the first step of TC-NER (Figure 2D). Importantly, all agents also caused a marked and specific increase of Pol II CTD Ser7^P^ (Figures 2D, S2D and S2E). These data demonstrate that Ser7 hyper-phosphorylation is a universal hallmark of elongation-stalled transcription, occurring in response to all tested obstacles that block the progression of elongating Pol II.

### Transcription stalling causes Ser7 hyper-phosphorylation *in trans*

To further understand stalling-induced Pol II Ser7 hyper-phosphorylation, we profiled the distribution of Ser7^P^ and total Pol II using dxChIP-seq, before and after UV irradiation. K1268R cells were used to prevent UV-induced RPB1 degradation and displayed near identical overall distribution of Ser7^P^ and total Pol II across gene classes as WT cells (Figures 1A and S2F). Exposure to UV caused a reduction of the transcription start site (TSS)-proximal total Pol II and Ser7^P^ signals across the genome (Figures 2E and S2G), which could reflect previously reported release of paused Pol II into elongation upon UV^52,53^.

Surprisingly, UV-induced Ser7 hyper-phosphorylation was detected in the early elongation zone, peaking 500 bp downstream of the TSS, as evidenced by increased Ser7^P^ signal (Figures 2E and 2F) and Ser7^P^ to total Pol II ratio (Figures 2E and 2G). If Ser7 hyper-phosphorylation occurred *in cis*, directly on elongating Pol II stalled at DNA damages, intuitively one would expect a different distribution of Ser7^P^ signal in dxChIP-seq, with Ser7^P^ to total Pol II ratio peaking further inside genes rather than in the early elongation zone (Figure S2H). Thus, the observed location of Ser7 hyper-phosphorylation suggests an *in trans* mechanism, whereby stalling of elongating Pol II in gene bodies triggers Ser7 hyper-phosphorylation on Pol II molecules at a remote part of the gene – at gene beginnings.

To test this possibility we used mathematical modelling^54^, to simulate Pol II dynamics after UV irradiation, in a transcription system containing one 100 kb-long gene and a pool of 250 Pol II molecules. Previously determined parameters describing Pol II behavior (such as initiation frequency, pausing duration, pause-termination rate, elongation speed, processivity, stalling halflife) and DNA damage (damage frequency, repair half-life) were used^25,55^ (for a full list of parameters see Table S1). This approach allowed us to simulate and contrast two different scenarios for Ser7 hyper-phosphorylation: 1) *in cis*, with Ser7 phosphorylated directly at polymerases stalled at DNA damage sites, and 2) *in trans*, where after the first Pol II encounters DNA damage and stalls, it sends a “signal” to the gene beginning to phosphorylate all polymerases passing through a “gate” located closely after the promoter-proximal pausing site, 200 bp from the TSS (Figure 2H). As the model operates in 100 bp increments along the gene, the “gate” for the *in trans* model was chosen to occur at the first increment following the promoter-proximal pausing site. However, the exact position of the “gate” (e.g. 200 bp vs 500 bp from the TSS) is not expected to substantially alter the results of the *in trans* model.

Remarkably, the *in trans* model recapitulated the results observed in the dxChIP-seq experiment, with Ser7^P^ signal peaking in the early elongation zone (Figures 2F-2H). As intuitively expected (Figure S2H), the *in cis* model simulated a relatively uniform and spread-out Ser7^P^, accumulating in the gene bodies, which does not resemble dxChIP-seq data (Figures 2F-2H). Interestingly, the *in trans* model also predicted a decline of Ser7^P^ signal compared to total Pol II below zero further inside gene bodies (Figure 2H). This prompted us to re-examine the dxChIPseq data beyond the 5 kb mark (Figure 2G) – indeed, the decline of Ser7^P^ predicted by the simulation can be observed in dxChIP-seq data (Figures 2H and 2I), albeit closer to the gene beginning (at ∼5 kb in dxChIP-seq data and at ∼20 kb in the simulation). A likely reason for this shift is the difference in Pol II speed along the gene, which was not simulated in the model for simplicity. Namely, this region (5-20 kb from the TSS) is exactly the place where Pol II transitions from slow to fast elongation^56–60^, which would explain the more rapid decline of signal in dxChIPseq data^55^ compared to the simulation, where Pol II speed was uniform along the gene.

Together, these results indicate that transcription stalling triggers Ser7 hyper- phosphorylation *in trans*, on polymerases leaving the pausing site and entering the early elongation zone. In support of this, using damaged “mini-gene” DNA templates for *in vitro* transcription with nuclear extracts is not sufficient to trigger Ser7 hyper-phosphorylation above the level seen with undamaged DNA templates. This indicates that a *trans*, signaling and/or topological component, is missing in the reaction (Figure S2I).

### Ser7^P^ constitutes a pathway separate from TC-NER and the “last resort” mechanisms

Upon transcription stalling, Ser7 hyper-phosphorylation occurs in concert with TC-NER and the “last resort” pathway – we therefore determined if it is a component of these mechanisms or constitutes a separate pathway altogether.

Eliminating TC-NER (by knocking-out CSB) and the “last resort” pathway (by RPB1 K1268R mutation) did not prevent Ser7 hyper-phosphorylation after UV (Figure 3A). In parallel, Ser7^P^ was not required to trigger TC-NER and the “last resort”, since FLAG-tagged S7A mutant Pol II can still recruit CSB (Figure 3B) and become poly-ubiquitylated upon stalling (Figure 3C). Thus, Ser7 hyper-phosphorylation, TC-NER and the “last resort” are triggered independently of each other upon transcription stalling. Of note, accumulation of Ser7^P^ seems to occur with a delay compared to TC-NER and Pol II ubiquitylation – it is most pronounced at timepoints when degradation of RPB1 is exhaustive, such as 3-6 h after UV treatment (Figures 2D, 3A and S2A). Due to this, for experiments in WT and S7A SWO cells that do not use proteasome inhibitors, we used short timepoints after UV, before RPB1 degradation takes place which is also before Ser7^P^ peaks (Figures 2B and 2C). Interestingly, CPT causes substantially less RPB1 degradation compared to other stalling agents (Figure S2D), and in the CPT condition Ser7 hyperphosphorylation can be observed in WT cells at earlier timepoints and without the use of proteasome inhibition or K1268R mutation (Figures S2D and S2E). These observations suggest that Ser7^P^ and RPB1 degradation may be used in complementary ways to deal with transcription stalling.

**Figure 3.**
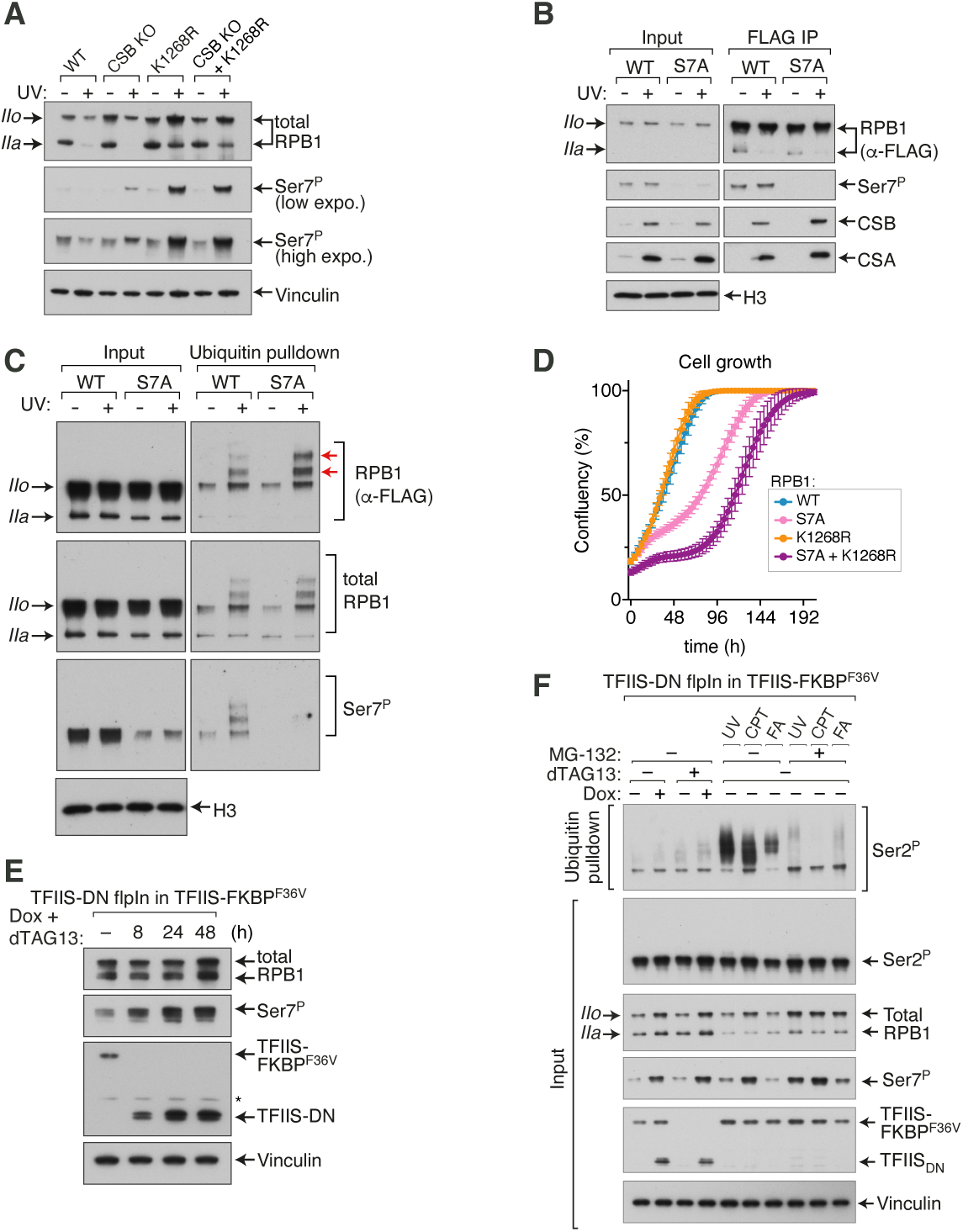
Ser7 hyper-phosphorylation is a separate pathway from TC-NER and the “last resort”, and is triggered by Pol II backtracking. **(A)** Western blot detecting total and Ser7^P^ RPB1, in WT, *CSB* KO, RPB1 K1268R and double mutant cells, before and 3h after UV treatment (20 J/m^2^). **(B)** Immunoprecipitation of RPB1-FLAG from chromatin, before and 45 min post-UV (20 J/m^2^), in WT and S7A switchover cells (transgenic RPB1-FLAG was induced ectopically with Dox but endogenous RPB1 was not eliminated with siRNA due to the scale of the experiment). Western blot detects ectopically expressed WT and S7A RPB1 using anti-FLAG antibody, as well as Ser7^P^ (which represents a combination of endogenous and ectopically expressed RPB1), CSB and CSA. **(C)** Ubiquitin pulldown from chromatin of WT and S7A switchover cells (endogenous RPB1 present, as in (b)), before and 25 min after UV treatment (20 J/m^2^). FLAG antibody specifically detects ectopically expressed WT and S7A RPB1, while D8L4Y and Ser7^P^ antibodies detect a combination of endogenous and ectopically expressed RPB1. Red arrows indicate polyubiquitylated RPB1 species. **(D)** Growth of WT, S7A, K1268R and double mutant (S7A + K1268R) switchover cells, monitored by Incucyte. A representative of biological triplicate experiment is shown, each with 6 technical replicate wells. Data shown as mean +/– standard error of imaging. Graph derived from the same experiment as Figure 1e. **(E)** Western blot detecting total RPB1 and Ser7^P^, in TFIIS_DN_ switchover cells. Degradation of endogenous, degron-tagged TFIIS-FKBP^F36V^ was induced by adding dTAG13 (100 nM), and expression of dominant negative TFIIS mutant (TFIIS_DN_) was induced with doxycycline. **(F)** Ubiquitin pulldown assay followed by Western blot detecting Ser2^P^ RPB1 (elongating Pol II) in TFIIS_DN_ switchover cells. Degradation of endogenous, degron-tagged TFIIS-FKBP^F36V^ was induced by adding dTAG13 (100 nM), and expression of dominant negative TFIIS mutant (TFIIS_DN_) was induced with doxycycline for 8 hours. For comparison, stalling was induced with UV, CPT and formaldehyde in cells expressing endogenous, wild-type TFIIS. In inputs, total, Ser2^P^ and Ser7^P^ are also detected.

Importantly, abolishing TC-NER and the “last resort” enhances Ser7 hyper-phosphorylation (Figure 3A). In *CSB* KO cells, Ser7 hyper-phosphorylation is evident even despite marked RPB1 degradation (Figure 3A). Likewise, abolishing Ser7^P^ enhances the recruitment of CSB and poly-ubiquitylation of stalled Pol II to some degree (Figures 3B and 3C). It has also been reported that abolishing TC-NER results in upregulation of the “last resort” activity^25,61^. Therefore, these three pathways might compensate for each other’s absence – when one is inhibited, the other two experience compensatory upregulation.

Next, we tested if Ser7^P^ could inhibit TC-NER and the “last resort” – if so, we would expect Ser7^P^ signal to be absent on CSB-bound or poly-ubiquitylated Pol II, respectively. The presence of Ser7^P^ on Pol II did not interfere with CSB recruitment, as Ser7^P^ can be detected on Pol II immunoprecipitated with CSB (Figure S3A). Similarly, Ser7^P^ is present on poly-ubiquitylated RPB1 after UV (Figure S3B). This was the case in both WT and *CSA* K.O. cells, suggesting that Ser7^P^ does not directly interfere with RPB1 poly-ubiquitylation, by either CSA or the “last resort” E3 ubiquitin ligase(s) (Figure S3C). Hence, Pol II Ser7 hyper-phosphorylation can simultaneously occur on the same Pol II molecules targeted by TC-NER or the “last resort”.

Interestingly, while K1268R mutation has no phenotype on its own in untreated cells, mutating this residue in combination with S7A (in the switchover model system) results in a synthetic growth impairment phenotype (Figure 3D). This indicates that Ser7^P^ and K1268ubiquitylation might be partially functionally redundant, possibly to deal with endogenous levels of Pol II stalling.

### Pol II backtracking triggers the Ser7^P^ pathway

A wide range of obstacles to elongating Pol II induce Ser7 hyper-phosphorylation, despite their fundamentally different physical properties (Figures 2A, 2D, S2A, S2D and S2E). We reasoned these different perturbations must have a common molecular basis – which is neither TC-NER nor the “last resort”, but some other event that is universally triggered by transcription stalling. We hypothesized that this could be Pol II backtracking. Elongating Pol II frequently backtracks, resulting in protrusion of RNA 3’end through the Pol II NTP pore^62–65^. At this point, TFIIS is recruited and stimulates hydrolysis of protruding RNA, allowing transcription to resume^66–68^. Obstacles in front of elongating Pol II dramatically increase the frequency of backtracking^15^, thus it is possible that backtracked Pol II might be a universal signal triggering Ser7 hyperphosphorylation. To test this, we induced expression of a dominant-negative mutant of TFIIS (TFIIS_DN_), which traps Pol II in the backtracked state but does not allow RNA hydrolysis (Figure S3D)^69^. We also tagged endogenous TFIIS with a degron tag (FKBP^F36V^)^70^ in these cells, enabling us to remove it. Within a few hours of TFIIS_DN_ expression, Ser7 became hyper-phosphorylated while phosphorylation at other CTD residues remained relatively unchanged (Figures 3E and S3E). This occurred irrespective of the presence or absence of endogenous, WT TFIIS (both in -dTAG and +dTAG conditions).

Trapping Pol II in the backtracked state with TFIIS_DN_ caused a similar extent of Ser7 hyperphosphorylation as the stalling agents UV, CPT and FA, when degradation of RPB1 was prevented in UV, CPT and FA conditions using MG-132 (Figures 3F and S3F). In contrast, TFIIS_DN_ caused very little to no poly-ubiquitylation of elongating, Ser2^P^ Pol II (Figure 3F), and no degradation of RPB1 (Figures S3F and S3G), even when TFIIS_DN_ was expressed for two days (Figures 3E and S3G). TFIIS_DN_ expression also caused substantially less CSB recruitment to Pol II than the stalling agents UV, CPT and FA (Figure S3F). This further shows that Ser7 hyper-phosphorylation is uncoupled from TC-NER and the “last resort” pathways, which do not seem to be efficiently activated by backtracked Pol II bound by TFIIS. In support of our data, recent cryo-electron microscopy (cryo-EM) structures show that the TC-NER factor UVSSA binds to the same regions of Pol II as TFIIS^71^. Thus, they compete and are unlikely to bind Pol II at the same time.

Surprisingly, MG-132 treatment not only prevented degradation of RPB1 upon UV, CPT and FA treatments, but also suppressed ubiquitylation of elongation-stalled, Ser2^P^ Pol II (Figure 3F). Although the underlying cause for this observation is unclear, it again indicates that Ser7 hyper-phosphorylation is activated when ubiquitylation and degradation of stalled Pol II is impaired, in agreement with results above using RPB1 K1268R mutants.

These findings suggest that persistence of stalled Pol II increases the likelihood of Pol II backtracking, which in turn triggers hyper-phosphorylation of Pol II Ser7. Notably, overexpression of TFIIS_DN_ causes most prominent Pol II backtracking in the first 2 kb from the TSS^63^ – the same area of the gene where we observe Ser7 hyper-phosphorylation upon UV irradiation (Figures 2E-2I).

### Ser7^P^ is required for transcription recovery after stalling

We next asked what the function of UV-induced Ser7 hyper-phosphorylation is. UV-lesions trigger global transcription changes, occurring in three phases^25,72,73^. First, sites of DNA damage act as physical barriers to elongating Pol II. Second, the “last resort” depletes cellular Pol II pool, exhausting the amount of Pol II available for initiation. The first two phases are often referred to as “a global transcription shutdown”^15^. Third, transcription starts to recover as DNA damages are repaired, allowing elongating Pol II to reach gene ends. During this time, the activity of the “last resort” also diminishes, allowing for subsequent recovery of the RPB1 protein level, which typically occurs with a delay, around 24 h after exposure to UV^25^. We tracked transcription shutdown and recovery using TT_chem_-seq. WT cells exhibited the expected transcription shutdown, with the first phase – impairment of Pol II elongation – observed 45 minutes after UV (Figures 4A and 4B). The second phase, initiation shutdown due to “last resort” RPB1 degradation, can be observed in WT cells 3 hours after UV (Figures 4A and 4B). These effects agree with previous reports^25,72,73^. In contrast, S7A cells displayed both elongation impairment and initiation shutdown already 45 minutes post-UV, and nascent RNA signal was further lost 3 h post-UV (Figures 4A and 4B). This suggested that excessive UV-induced degradation of RPB1 may occur when Ser7^P^ is abolished, which was indeed confirmed by western blot (Figure 4C).

**Figure 4.**
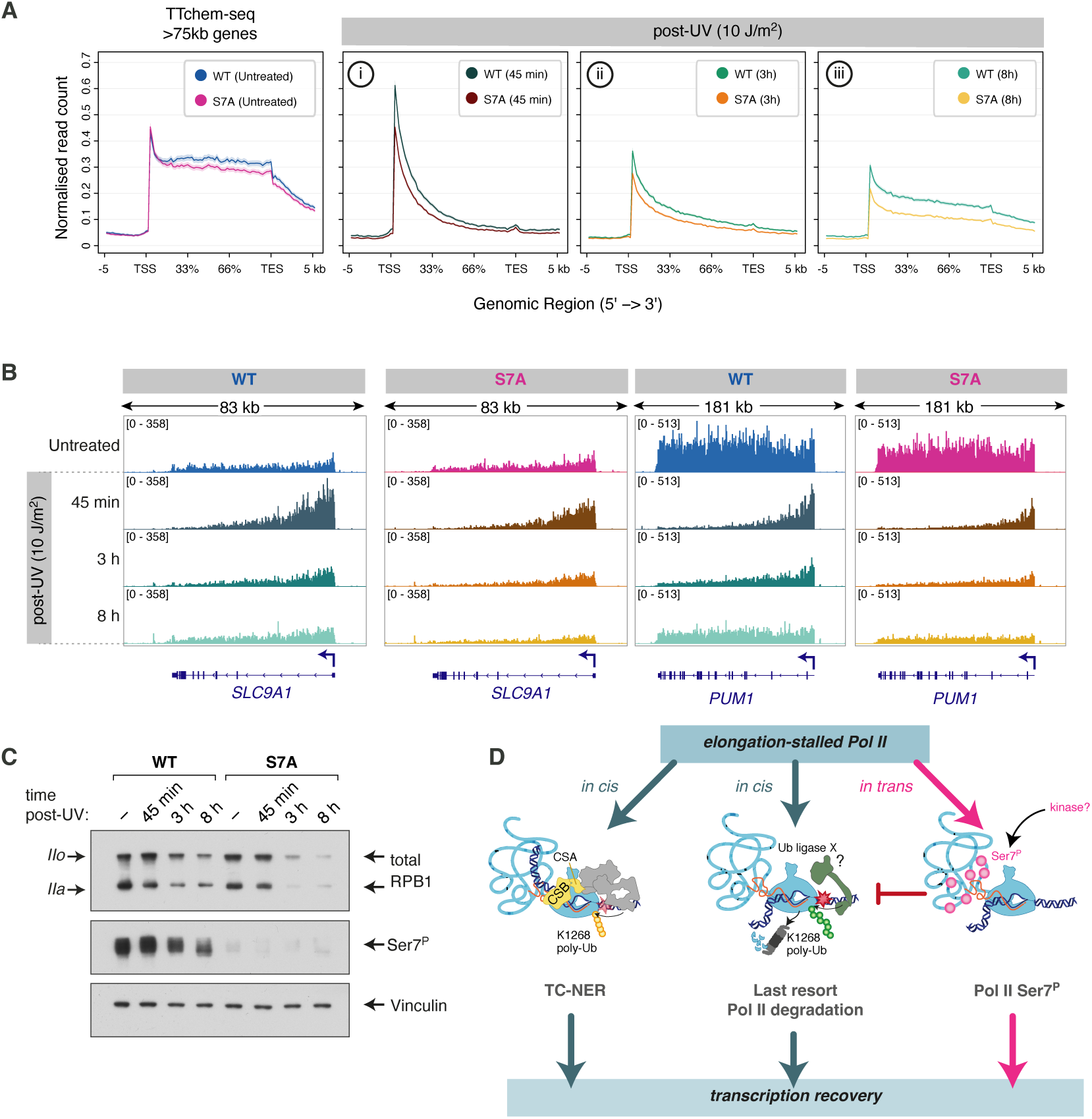
Ser7^P^ is required for transcription recovery upon DNA damage. **(A)** Metagene plot of TT_chem_-seq data, showing nascent RNA production on long genes (>75 kb), for WT and S7A switchover cells, before and at varying time-points after UV (10 J/m^2^). Timepoints capture the first and second stage of transcription shutdown (45 min and 3h, respectively) and the first stage of transcription recovery (8h). **(B)** Example of nascent RNA signal (TT_chem_-seq data) on two long genes (*SLC9A1* and *PUM1*), for WT and S7A switchover cells, before and at varying time-points after UV (10 J/m^2^). **(C)** Western blot detecting total and Ser7^P^ RPB1 in WT and S7A switchover cells, before and at varying time-points after UV (10 J/m^2^). **(D)** Schematic illustrating the function and relationship between pathways triggered in response to transcription stalling.

Furthermore, onset of transcription recovery was evident 8 hours post-UV in WT cells (phase 3), but was impaired in S7A cells (Figures 4A and 4B). Interestingly, similarly to WT cells, Pol II could reach ends of genes in S7A cells 8 hours post-UV, but the overall level of transcription was lower throughout the gene (Figures 4A, 4B and S4). This indicates that DNA damages have been repaired both in WT and S7A mutant cells, suggesting that TC-NER is proficient. Instead, transcription recovery is impaired due to the excessive degradation of RPB1 protein in S7A cells, which can be observed by western blot (Figure 4C). In agreement, RPB1 poly-ubiquitylation after UV is enhanced in S7A mutants compared to the WT (Figure 3C). Therefore, Ser7 hyperphosphorylation protects stalled Pol II from excessive destruction by the “last resort” pathway. This likely explains the apparent complementary behavior of Ser7^P^ and RPB1 degradation observed throughout the study (Figures 2A, S2A, S2D, S2E, 3A, 3F, S3E-S3G).

Taken together, like TC-NER and the “last resort”, Ser7^P^ is required for transcription recovery after DNA damage, but unlike TC-NER and the “last resort” which operate *in cis* – directly on stalled Pol II molecules^24,27,28^, Ser7 hyper-phosphorylation occurs *in trans*, in the early elongation zone. Notably, while all three pathways are required for transcription recovery after DNA damage, only Ser7^P^ is also required for normal cellular function in the baseline state (Figures 1E-1H). Abolishing TC-NER or the “last resort” has no adverse effects on cell growth or transcription in untreated condition^25^. This suggests that Ser7^P^ might be a primary pathway to deal with endogenous obstacles to elongating Pol II, while all three pathways may be necessary when the burden of roadblocks on genes is increased.

### GSK3 is a Pol II CTD Ser7 kinase

We next sought to identify Ser7 kinase. Interestingly, Glycogen Synthase Kinase 3 (GSK3) has recently been implicated in the phosphorylation of Pol II upon UV irradiation, however the target residue on the Pol II CTD remained unclear^74^. To determine which Pol II CTD residue or residues GSK3 phosphorylates, we treated K1268R cells with a specific GSK3 inhibitor CHIR-99021, induced transcription stalling with UV irradiation, and assessed phosphorylation at Pol II CTD Ser2, Ser5 and Ser7. GSK3 inhibition in cells dramatically reduced the level of Ser7^P^, both in untreated and UV-irradiated cells, while also causing a minor reduction of Ser2^P^ (Figure 5A). To test this more directly, we performed *in vitro* transcription assays with nuclear extracts, whereby GSK3 was inhibited with CHIR-99021 thirty seconds before transcription (and phosphorylation) was initiated by the addition of NTPs. In this assay, inhibition of GSK3 completely abolished Ser7^P^ and also reduced Ser2^P^ (Figure 5B). The latter effect is comparable with the one seen with CDK9 inhibition by 5,6-Dichloro-1-β-D-ribofuranosylbenzimidazole (DRB) (Figures 5B and S5A). Considering that Ser7^P^ has been proposed to prime the CTD for Ser2-phosphorylation by CDK9^75^, it is unclear if the effect of GSK3 on Ser2^P^ in cells and in nuclear extracts is direct or not.

**Figure 5.**
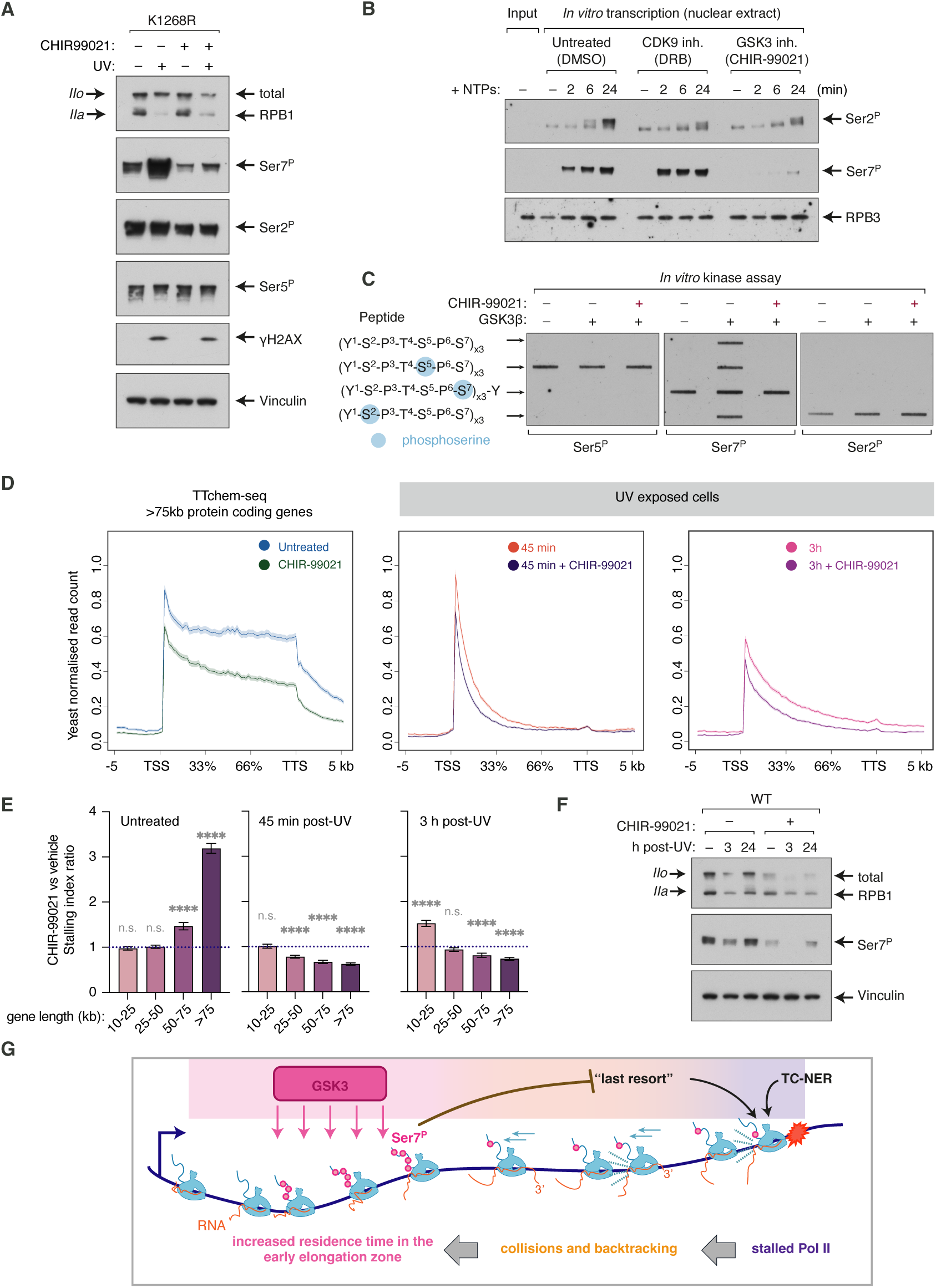
GSK3 is a Pol II Ser7 kinase. **(A)** Western blot detecting total RPB1 and its phospho-forms, upon treatment with UV (6 h post-UV, 30 J/m^2^), and inhibition of GSK3 (20 µM CHIR-99021, cells pre-treated for 3 h), in RPB1 K1268R cells. **(B)** *In vitro* transcription assay with nuclear extracts, followed by Western blot detecting Ser7^P^ and Ser2^P^ RPB1. Inhibitors of CDK9 (DRB, 100 µM) and GSK3 (CHIR-99021, 10 µM) were added 30 s before initiating transcription and phosphorylation by adding NTPs. RPB3 subunit of Pol II is used as a loading control. **(C)** *In vitro* kinase assay with recombinant GSK3 and synthetic Pol II CTD tri-peptides. After incubation with GSK3 and ATP (with or without GSK3 inhibitor CHIR-99021), the peptides were immobilized on a membrane and phosphorylation at different residues (Ser2, Ser5 and Ser7) was detected using immunoblotting. **(D)** Metagene plot of TT_chem_-seq data, showing nascent RNA production on long genes (>75 kb), in WT cells with or without CHIR-99021 treatment, before and at varying time-points after UV (15 J/m^2^). **(E)** Comparison of stalling indices derived from TT_chem_-seq data, in CHIR-99021-treated vs untreated WT cells, for different gene length categories. Values above 1 indicate more Pol II stalling in CHIR-99021-treated compared to untreated condition (indicated with blue dashed line). Two-tailed t-test comparing CHIR-99021 vs vehicle treated samples (n.s. = not significant, **** = p<0.0001). **(F)** Western blot detecting total and Ser7^P^ RPB1, 3 h and 24 h after treatment with UV (20 J/m^2^), and inhibition of GSK3 (20 µM CHIR-99021, cells pre-treated for 3 hours), in WT cells. **(G)** A simplified model for how stalling of elongating Pol II in gene bodies could activate the GSK3-Ser7^P^ pathway at a distal part of the gene, *in trans*.

Following this, we performed an *in vitro* kinase assay using purified GSK3β and triheptamer CTD peptides chemically synthesized with pre-existing phosphorylation at Ser2, Ser5 and Ser7. In this assay, GSK3β exclusively phosphorylated Ser7, irrespective of the pre-existing phosphorylation status of the tri-peptide substrates (Figure 5C). Together, these results show that GSK3 phosphorylates Pol II CTD Ser7 *in vivo* and *in vitro*, and that it may also be able to phosphorylate Ser2 in certain contexts.

Next, we probed the interaction between Pol II and GSK3 by immunoprecipitation of GSK3β. GSK3β interacts with Pol II in untreated cells, and this interaction is further enhanced by UV irradiation (Figure S5B).

Inhibition of GSK3 phenocopied the effects of S7A mutation on nascent RNA transcription both before and after UV, as determined by TT_chem_-seq. In the baseline state, CHIR-99021 impaired elongation, particularly on long genes (Figures 5D and 5E). Like S7A, CHIR-99021 treatment exacerbated transcription shutdown after UV, by causing an overall reduction in nascent RNA production throughout genes (Figures 5D and S5C), most likely due to depletion of RPB1 protein (Figure 5F). Peculiarly, both GSK3 inhibition and S7A mutation partially mitigate elongation-inhibitory effects of UV irradiation genome-wide (Figures S4 and 5E, right), despite depleting RPB1 protein (Figures 4C and 5F). In agreement, the Kornblihtt laboratory has recently reported that CHIR-99021 treatment counteracts UV-induced inhibition of transcription elongation on a model gene^74^. These findings suggest that GSK3 → Pol II Ser7^P^ pathway stimulates transcription elongation in unperturbed cells, yet slows it down when the burden of roadblock on genes is increased, such as after UV irradiation.

Interestingly, CHIR-99021 treatment alone (in otherwise untreated cells) additionally caused a reduction of RPB1 protein level (Figure 5F) and nascent RNA production globally (Figures 5D and S5C), while S7A mutation does not have these effects in untreated cells. These differences may arise due to the inability of cells to compensate for the rapid reduction in Ser7^P^ caused by acute GSK3 inhibition, whereas they may be capable of activating compensating mechanisms upon chronic Ser7^P^ depletion (caused by S7A mutation). Alternatively, given that GSK3 targets over 100 substrates in the cell^76^, it is possible that the effect of CHIR-99021 on RPB1 level and global nascent RNA production in the baseline state is mediated through additional transcriptional effectors GSK3 may phosphorylate.

Together, these data indicate that GSK3 is a Pol II CTD Ser7 kinase. GSK3 phosphorylates Pol II at Ser7 residues both in the baseline state and in response to transcriptional stalling.

## Discussion

Here, we uncover the role of Ser7^P^ in Pol II transcription and define the mechanism of its biogenesis in human cells. While transcription initiation and pause-release are relatively wellunderstood stages of the transcription cycle, recent studies and the data presented here clearly indicate that additional downstream steps of regulation exist^42,58,77–79^. Deposition of Ser7^P^ in the early elongation zone and its role in supporting elongation suggest that specific molecular transactions occur in this area of genes to prepare Pol II for efficient elongation. Using PRO-IPseq, Vihervaara *et al*. observed that the higher Ser7^P^ in the early elongation zone, the more processive Pol II was on a given gene^42^. In agreement, our data demonstrate that abolishing Ser7^P^ impairs elongation efficiency and causes premature termination, particularly on long genes. This supports a model whereby Ser7^P^ stimulates assembly of elongation complexes in the early elongation zone to enable processive transcription throughout the rest of the gene. To understand the mechanisms behind this, in the future it will be important to identify the factors that “read” the Ser7^P^ mark.

Interestingly, Integrator has been proposed to bind Ser7^P^ and mediate transcription termination at snRNA genes^14,80,81^. However, recent cryo-EM structures show that phosphorylation at Ser7 is incompatible with Integrator binding to Pol II CTD and might in fact displace it from Pol II rather than recruit it^82^. Nonetheless, Integrator is an important regulator of transcription in the early elongation zone^77,78,83–92^, thus it will be relevant to determine if and how it may be connected to Ser7^P^ function on protein-coding genes. Regardless, it is likely that not one, but multiple proteins bind Ser7^P^ to mediate its effects, as is common with other Pol II CTD marks^3,5,41,93,94^.

Other Pol II CTD phospho-marks (Tyr1^P^, Ser2^P^, Thr4^P^ and Ser5^P^) also have important and distinct functions in regulating transcription, and their levels are thought to be dynamic but constitutive^2–6^. Remarkably, we observe Ser7 is dramatically hyper-phosphorylated in response to transcription stalling. Peculiarly, experiments and mathematical modelling show this Ser7 hyperphosphorylation occurs *in trans* – in the early elongation zone – and not *in cis* – directly on stalled Pol II. This raises a question: how is the signal propagated from the site of stalling to the early elongation zone? Since all types of transcription-stalling obstacles we tested in this study caused Ser7 hyper-phosphorylation, the underlying “trigger” must be a common molecular event, and our data suggest that Pol II backtracking is involved. From this point, we envisage two possibilities: there could be unknown signaling factors that sense stalled and/or backtracked Pol II in gene bodies and cause Ser7 phosphorylation *in trans*, in the early elongation zone. Alternatively, *in trans* Ser7 hyper-phosphorylation could be caused by changed dynamics of the entire transcription system when there are obstacles on genes. Namely, when one Pol II molecule stalls on a gene, Pol II molecules travelling behind it will soon encounter it, forming queues of collided polymerases that are prone to backtracking. This causes crowding of Pol II molecules between the site of damage and the gene beginning (evident in dxChIP-seq profiles) which is likely to overall slowdown the “traffic” of Pol II on the gene. As a result, Pol II would simply have an increased residence time in the early elongation zone where Ser7 is normally phosphorylated, enhancing the density of Ser7 phosphorylation per Pol II molecule (Figure 5G). This scenario follows Occam’s razor principle, potentially explaining Ser7 hyper-phosphorylation *in trans* without assuming the existence of additional signaling molecules. Moreover, it is possible that backtracked Pol II itself is a favorable substrate for GSK3 to phosphorylate Ser7 – it has been shown that most extensive Pol II backtracking occurs exactly in the early elongation zone^62,63^, which is likely due to reduced Pol II speed in this area of genes^42,56–60^.

Multiple kinases phosphorylate Pol II throughout the transcription cycle, and seem to act in a defined, successive order^3–6^. We reveal that GSK3 phosphorylates Pol II CTD Ser7, both in the baseline state and upon induction of transcription stalling. How GSK3 activity is coordinated with other transcriptional kinases remains to be determined. In our assays, GSK3 inhibition also reduced Ser2^P^, but this effect may be indirect, as Pol II CTD harboring Ser7^P^ has been proposed as a favorable substrate for CDK9 to phosphorylate Ser2^75^. Nonetheless, Ser7^P^ density peaks just after Pol II leaves the pausing site, suggesting that GSK3 acts on Pol II after CDK7 (which phosphorylates Ser5) but before or in concert with CDK9. Furthermore, interaction between transcriptional kinases likely extends beyond coordinating their sequential action. GSK3 prefers substrates that harbor priming pre-phosphorylation at a defined distance from its own target residue^95^. Indeed, examining the Pol II CTD motif with a recently developed computational prediction tool, The Kinase Library^96^, returns GSK3b and GSK3a as top hits for Pol II CTD Ser7 kinases with very high (>99.9%) confidence – but only if the downstream Thr4 residue is prephosphorylated as a priming event. In the future, it will be important to test this prediction *in vivo*, and to determine if there could be additional Ser7 kinases in different cellular contexts.

Interestingly, GSK3 has recently been proposed to stimulate degradation of promoterproximal Ser5^P^ Pol II after UV, but independently of the “last resort”^97^. We and others subsequently demonstrated that promoter-proximal Ser5^P^ Pol II is a substrate of CRL3^ARMC5^ ubiquitin ligase, which monitors the quality and quantity of Pol II molecules before they are released into elongation^77,78^. We also found that this process is entirely separate from the “last resort” RPB1 K1268-ubiquitylation and degradation that operates on elongating Pol II^77^. It is therefore possible that GSK3 not only phosphorylates Pol II at Ser7 in the early elongation zone, but also regulates the activity of CRL3^ARMC5^. It is plausible that upon stalling of Pol II in gene bodies, reducing the number of Pol II molecules that are released into elongation would be beneficial, to avoid exacerbation of Pol II collisions, crowding and stalling throughout the gene, and GSK3 ® CRL3^ARMC5^ axis may fulfil this function. This may also provide an alternative or additional explanation for the reduction in Pol II occupancy at promoter-proximal regions upon UV – this could be the result of enhanced pause-release rate as previously proposed^52,53^, but also the result of Pol II degradation by CRL3^ARMC5^, or possibly even increased pause-termination rate. Going forward, it will be important to determine if and how GSK3-Pol II Ser7^P^ and CRL3^ARMC5^, two newly discovered transcriptional mechanisms, are connected.

Pol II elongation remains the least understood step of transcription, with only two mechanisms known so far to resolve stalled elongating Pol II complexes: TC-NER and the “last resort” pathway^15^. This work demonstrates that GSK3-Pol II Ser7^P^ is a separate, third pathway triggered in response to stalling of elongating Pol II. Like TC-NER and the “last resort”, GSK3Pol II Ser7^P^ is required for transcription recovery after DNA damage, but unlike TC-NER and the “last resort”, GSK3-Pol II Ser7^P^ acts *in trans*, not *in cis*. We propose that crowding of Pol II molecules on the gene (resulting from stalling and backtracking of elongating Pol II) provides a simple “sensor” of the burden of obstacles on genes – extending the residence time of Pol II in the early elongation zone where it becomes hyper-phosphorylated at Ser7. This Ser7 hyperphosphorylation is functional: it “warns” polymerases in the early elongation zone that there are roadblocks ahead. In this model, increased density of Ser7^P^ prepares these polymerases to face obstacles further along the gene, likely by recruiting unknown factor(s) that will protect elongating Pol II from excessive degradation – restraining the “last resort” from exhausting the cellular Pol II pool needed for transcription recovery (Figure 5G).

## RESOURCE AVAILABILITY

### Lead contact

Further information and requests for resources and reagents should be directed to and will be fulfilled by the lead contact, Ana Tufegdžić Vidaković.

### Materials availability

Materials generated in this study will be made available on request, but we may require a completed Materials Transfer Agreement if there is potential for commercial application.

### Data and code availability

#### Data

All data are available in the main text or supplementary materials. TT_chem_-seq and dxChIP-seq data are deposited to GEO under the accession number GSE304829. Reagents used in this study are available from A.T.V. upon request.

#### Code

All tools and software used in the analyses are publicly available, and details are described in STAR Methods. Custom code is available at https://github.com/shutongye/bcs_Pol_II_phosphorylation.

## Acknowledgements

We thank Julian E. Sale. Lori Passmore, Mariann Bienz, Kelly Nguyen and Diana Arseni for critical reading of the manuscript. We thank members of the A.T.V. lab for productive discussions and the MRC LMB Facilities for supporting our work (Flow cytometry, Mechanical and Electronics Workshops, Media kitchen, Scientific Computing). We are grateful to Mariann Bienz and Tiemei Li for a kind gift of purified GSK3b, and to Jesper Q. Svejstrup for a kind gift of pcDNA-TCEA1mut plasmid, *CSB* K.O. and RPB1 K1268R cell lines and Pol II CTD peptides. A.T.V. is supported by a core grant to the LMB from the Medical Research Council (Ref. MC_UP_1201/28) and acquired preliminary data with support of Francis Crick Institute funding (FC001166). I.S. is supported by a César Milstein Studentship from the Darwin Trust of Edinburgh.

## Author contributions

I.S. performed experiments, analyzed data, and wrote the manuscript. Y.B. performed experiments, analyzed data, and wrote the manuscript. T.T. performed experiments and analyzed data. A.C. analyzed data. S.Y. performed mathematical modeling. S.W. analyzed data. R.C. performed experiments. A.E. developed methodology. D.C. developed methodology. M.B. performed mathematical modelling and provided mentoring and supervision. A.T.V. conceived the project, acquired funding, provided mentoring and supervision, performed experiments, analyzed data and wrote the manuscript.

## STAR Methods

### Plasmids and oligonucleotides

All plasmids and oligonucleotides are listed in Supplementary Table S2.

### Cell lines and culture conditions

Cell lines used in this study are listed in Supplementary Table S3. All cell lines were cultured in supplemented high glucose DMEM (Gibco) with 10% fetal bovine serum (FBS, Gibco), 100 U/mL penicillin, 100 μg/mL streptomycin at 37 °C with 5% CO_2_. Switchover (SWO) cell lines were cultured in media with 10% tetracycline-free FBS (Biosera) and additionally supplemented with 100 μg/mL Hygromycin B (Invitrogen) and 15 μg/mL Blasticidin S (Gibco). All cell lines used in this study are derived from Flp-In T-Rex HEK293 and are female. Cells were regularly tested for mycoplasma. Cells were not authenticated.

### Generation of cell lines

*CSA* K.O., *TFIIS* K.O. and the double mutant RPB1 K1268R + *CSB* K.O. were generated using CRISPR Cas9 nickase system, each locus targeted by two gRNAs to excise the region between them^98^. gRNAs were assembled into the pX461.

SWO cell line generation was performed as previously described^25^. Briefly, pFRT-TO backbone plasmid containing the RPB1 (*POLR2A*) coding region with 6 x His tag in the C-terminal region, and synonymous mutations providing resistance to two siRNAs targeting RPB1 (D011186-03-0020 and D-011186-05-0020, Dharmacon) were obtained by synthesis and cloning from Epoch Bioscience. S7A variant with serine to alanine (S→A) mutation in all instances of serine in the 7^th^ position of the CTD was also obtained. Plasmids containing FLAG-tagged RPB1 variants were constructed by site-directed mutagenesis from the above plasmids. Flp-In T-REx HEK293 SWO cell lines were generated by co-transfection with the above plasmids and pOG44 Flp-recombinase expression vector (Thermo Fisher Scientific) followed by selection with 100 μg/mL Hygromycin B and 15 μg/mL Blasticidin S. Single colonies were isolated and expanded. Expression of SWO RPB1 was induced by the addition of doxycycline and verified by western blot using antibodies against His or FLAG tag.

Knock-in cells with the FKBP^F36V^ tag on N-terminus of TFIIS were constructed by CRISPRCas9-nickase mediated genome editing of Flp-In T-REx HEK293 cell lines, with modifications to the previously described method^70^. In short, homology-directed repair donor plasmids were generated by Gibson assembly with approximately 1000-bp flanking homology repair fragments PCR amplified from human gDNA, and an FKBP^F36V^-PuroR cassette amplified from pCRIS-PITChv2-dTAG-Puro (BRD4). Mutations were introduced in the homology arms by site-directed mutagenesis to provide resistance to guide RNAs. Cells were co-transfected with the donor plasmids and two pX461 Cas9 gRNA plasmids (with two gRNAs targeting the insertion site designed to face outwards) with Lipofectamine 3000 (Thermo Fisher Scientific). Cells were reseeded in medium containing 1 µg/mL Puromycin and cultured for two weeks. Single clones were isolated and verified by genomic PCR and western blot. TFIIS_DN_ (TFIISD282A-E283A) was cloned into pFRT-TO plasmids using pcDNA4/TO-TFIISmut plasmids^64^ as a template for Gibson assembly. This plasmid was then used to generate switchover TFIIS_DN_ in TFIIS-FKBP^F36V^, *TFIIS* K.O. and WT backgrounds, using the same protocol as for RPB1 SWO cells above.

### Cell treatments

For TT_chem_-seq and cell growth assays, siRNA transfections were performed with Lipofectamine RNAiMax (Thermo Fisher) according to manufacturer instructions, with 40 nM final siRNA concentration. UV irradiation was performed using a custom-built UV conveyor belt and the given dose was determined using a UV-meter with a detector set at 254 nm^34^. Doxycycline (Dox) induction was carried out at 1µg/mL, unless otherwise stated. A full list of reagents is listed in Supplementary Table S4.

### Cell growth assays

One 6-well plate was seeded containing 6 × 10^5^ SWO cells. The following day, the RPB1 transgene was induced with Dox, and cells transfected with siRNAs targeting RPB1 (D-011186-03-0020 and D-011186-05-0020, Dharmacon) 6-10 hours post-induction with Dox, using Lipofectamine RNAiMax (Invitrogen), according to manufacturer’s instructions. 40 nM total siRNA concentration was used. The following day 5,000 or 10,000 SWO cells were seeded per well in poly-D-lysine (Merck) coated 96-well plates, 12-18 hours after siRNA transfections. Growth was monitored and recorded every 3 h using Incucyte (Sartorius). Data from one representative experiment of three biological replicates, each with 6 technical replicate wells per condition and 4 imaging areas per well, were used for plotting growth curves.

### Dsk2 ubiquitin pulldown assays

Preparation of Dsk2-conjugated beads and ubiquitin pulldown assays were performed as previously described in detail^34^. Briefly, cells were harvested by scraping down in PBS, spinning down at 300 rcf and removing the supernatant. Cell pellets were resuspended in TENT buffer (50 mM Tris-HCl, pH 7.4, 2 mM EDTA, 150 mM NaCl, 1% (v/v) Triton X-100) containing fresh protease inhibitors, phosphatase inhibitors and 2 mM of N-Ethylmaleimide (NEM). Samples were incubated on ice for 10 min, sonicated in a 4 °C water bath (Bioruptor) at high power, with 30 s ON and 30 s OFF pulses, for a total duration of 7 min, then centrifuged at 18,000 rcf for 7 min. Dsk2 beads were pre-washed in TENT buffer containing fresh protease inhibitors, phosphatase inhibitors and 2 mM NEM. A bead suspension of 0.2–0.4 mL (equivalent to 10–20 μL packed beads) was used to pull down ubiquitylated proteins from 1–2 mg of the whole cell protein extract. Samples were incubated on a turning wheel at 4 °C overnight. The beads were then washed twice with TENT buffer containing fresh protease inhibitors, phosphatase inhibitors and 2 mM NEM, and then once with PBS containing protease inhibitors, phosphatase inhibitors and 2 mM NEM. The samples were typically eluted from beads with 40 μL of Laemmli buffer containing DTT. Samples were vortexed briefly, boiled at 98 °C for 5 min, spun down and supernatants were saved and analyzed by western blot. For one experiment (Figure 3C), chromatin samples (obtained using chromatin fractionation as described in immunoprecipitation section) were used instead of whole cell lysates.

### Western blot analysis

For whole cell extracts, cell pellets were lysed in protein lysis buffer (20 mM Tris-HCl pH 7.5, 250 mM NaCl, 1 mM EDTA, 0.5% (v/v) NP-40, 10% (v/v) glycerol, supplemented with protease inhibitors, phosphatase inhibitors and 2 mM NEM) and sonicated in a 4 °C water bath sonicator (Bioruptor) at high power, with 30 s ON and 30 s OFF pulses, for a total duration of 7 min, then centrifuged 18,000 rcf for 7 min. Total protein concentration was normalized and run by SDSPAGE on 4%–12% or 4%–20% Tris-Glycine gels (Thermo Fisher Scientific) and transferred onto 0.45 µm nitrocellulose membranes (Cytiva) using wet transfer. Membranes were blocked in 5% (w/v) skimmed milk in PBST (PBS, 0.05% (v/v) Tween 20) for 1 hour at room temperature, and incubated with primary antibodies (see table below) in blocking buffer overnight at 4 °C. Membranes were washed 3 times in PBST, incubated with HRP-conjugated secondary antibodies (see table below) in blocking buffer for 1 hour at room temperature. Protein detection was carried out using SuperSignal West Pico PLUS Chemiluminescent Substrate (Thermo Fisher Scientific) or Radiance Plus Femtogram HRP substrate (Azure Biosystems) according to manufacturer’s instructions. Signals were visualized by exposing the membrane to X-ray films (Fujifilm) and developed using an Optimax X-ray film developer (Protec). The list of antibodies used for western blot is listed in Supplementary Table S5.

### Immunoprecipitation

Phosphatase inhibitors, protease inhibitors and NEM were added fresh to all buffers. Cells scraped in PBS from one 15 cm dish were used per condition. For chromatin extraction, frozen pellets were quickly defrosted at room temperature and transferred to ice. 2 pellet volumes of Hypotonic Buffer (10 mM HEPES-KOH, pH 7.5, 10 mM KCl, 1.5 mM MgCl_2_) were added and the cells were incubated on ice for 20 min. To release cytosol, 20 strokes with a loose pestle were applied to the samples in a dounce-homogenizer. To pellet nuclei, the samples were centrifuged at 1,000 rcf at 4 °C for 20 min. Supernatant (cytosolic fraction) was removed, and pellets were resuspended by pipetting in 2 original pellet volumes of Nucleoplasmic Extraction Buffer (20 mM HEPES-KOH, pH 7.5, 1.5 mM MgCl_2_, 10% (v/v) glycerol, 150 mM potassium acetate, 0.05% NP-40), and incubated on a turning wheel at 4 °C for 20 min. To separate nucleoplasmic proteins from chromatin, the samples were centrifuged at 20,000 g at 4 °C for 20 min. The supernatant (nucleoplasmic fraction) was removed and chromatin pellets were resuspended in 1 pellet volume of Chromatin Digestion Buffer (125 U/mL benzonase (BaseMuncher Endonuclease, Abcam) in 20 mM HEPES-KOH, pH 7.5, 1.5 mM MgCl_2_, 10% (v/v) glycerol, 150 mM NaCl, 0.05% (v/v) NP-40). The samples were incubated on ice for at least 30 min, then centrifuged at 20,000 g at 4 °C for 20 min. Supernatant (first chromatin fraction) was saved in a new tube. The remaining chromatin pellet was resuspended in 1 pellet volume of Chromatin-2 Buffer (20 mM HEPESKOH, pH 7.5, 1.5 mM MgCl_2_, 10% glycerol (v/v), 3 mM EDTA, 500 mM NaCl, 0.05% (v/v) NP40) and incubated on a turning wheel at 4 °C for 20 min. 2.3 pellet volumes of Dilution Buffer (20 mM HEPES-KOH, pH 7.5, 1.5 mM MgCl_2_, 10% (v/v) glycerol, 3 mM EDTA, 0.05% (v/v) NP-40) were added to each sample, and samples were then centrifuged at 20,000 rcf at 4 °C for 15 min. Supernatant (second chromatin fraction) was saved and pooled with the first chromatin fraction and used as the total chromatin fraction in the following procedures.

1–2 mg of chromatin fraction was used per IP (all samples were adjusted to the same volume, typically 800-950 μL). For immunoprecipitation of RPB1 with 4H8, CSB or GSK3β antibody, 100 μL of packed Protein G agarose or magnetic beads (Thermo Fisher Scientific) per sample were prepared, by washing twice in 0.05% (v/v) Tween 20 in PBS (PBST) and then coupling to 10–30 μg of antibody per sample, for 1 h on a turning wheel at room temperature. FLAG immunoprecipitations were done using Anti-FLAG M2 affinity gel (Sigma-Aldrich). The beads were washed twice in PBST and once in IP buffer (20 mM HEPES-KOH, pH 7.5, 1.5 mM MgCl_2_, 10% (v/v) glycerol, 150 mM NaCl, 0.05% (v/v) NP-40). The beads were then resuspended in IP buffer and added to the samples, to yield a 1 mL total reaction volume. The samples were incubated on a turning wheel at 4 °C for 3 hours. Supernatant (unbound fraction) was removed, and the beads were washed 3 times in IP buffer. To elute immunoprecipitated proteins, 100 μL of 2X Laemli buffer were added to the beads, the beads were briefly vortexed and boiled at 98 °C for 5 min. The beads were centrifuged at maximum speed for 2 min and the supernatant (elution) was transferred to a new tube for further analysis by western blot.

### RT-qPCR

For RT-qPCR, total RNA was extracted using miRNeasy kit (QIAGEN), following the instructions of the manufacturer including an on-column DNase treatment (QIAGEN). 1 μg of RNA per sample were first denatured at 65 °C for 5 min and snap-cooled on ice, then used for reverse transcription with TaqMan Reverse Transcription Reagents or Superscript IV Reverse Transcriptase (Thermo Fisher Scientific) using random hexamers. 3 μL of cDNA diluted with water (typically 1:3–1:5) were used per well for qPCR with iTaq Universal SYBR Green Supermix (BioRad), with 40 cycles of 15 s denaturation at 94 °C, 30 s annealing at 58 °C, and 30 s extensions at 72 °C. Primers amplifying mature GAPDH were used as normalization control. Primer sequences are listed in plasmids and oligonucleotides session. Data were analyzed using Excel and plotted in GraphPad Prism. A minimum of three biological replicates were analyzed, each in technical triplicates.

### TT_chem_-seq (nascent RNA-seq)

TT_chem_-seq was performed essentially as described, with minor modifications^43^. One 70% - 80% confluent 15cm dish was used per condition in CHIR099021 treated cells. For SWO cells, 2 wells of a 6-well plate were seeded, each containing 6 × 10^5^ cells. The next day, the transgenic RPB1 was induced with Dox, and cells transfected with siRNAs targeting RPB1 (D-01118603-0020 and D-011186-05-0020, Dharmacon) 6-10 hours post Dox induction, using Lipofectamine RNAiMax (Invitrogen) according to manufacturer’s instructions. 40 nM total siRNA concentration was used. The next day, cells of the same condition were re-seeded and combined into a single 10 cm dish, in media containing Dox. The following day, cells were treated with UV, and nascent RNA was *in vivo* labelled with a 1 mM 4-thiouridine (4sU) (Glentham Life Sciences) pulse for exactly 15 min. Labelling was stopped by TRIzol (Thermo Fisher Scientific) and RNA was extracted as described previously^43^.

As a control for sample preparation, 4-thiouracil (4TU)-labelled RNA derived from *S. cerevisiae* (strain BY4741, MATa, his3D1, leu2D0, met15D0, ura3D0) was spiked into each sample. *S. cerevisiae* cells were grown in YPD medium overnight, diluted to an OD_600_ of 0.1, then grown to mid-log phase (OD_600_ approximately of 0.8) and incubated with 5 mM 4TU (Sigma-Aldrich) for 6 min. Yeast metabolizes 4TU into 4sU, which is incorporated into nascent RNA. Total yeast RNA was extracted using PureLink RNA Mini kit (Thermo Fisher Scientific) following the protocol.

For purification of 4sU labelled RNA, 100 μg of human 4sU-labelled RNA was spiked-in with 1 μg of 4sU-labelled *S. cerevisiae* RNA. The 101 μg of RNA (in a total volume of 100 μL) were fragmented by addition of 20 μL freshly made 1 M NaOH and incubated on ice for 20 min. Fragmentation was stopped by addition of 80 μL 1 M Tris-HCl, pH 6.8 and the samples were cleaned up twice with Micro Bio-Spin P-30 Gel Columns (BioRad) by adding 200 µL of RNA solution per column. The biotinylation of 4sU residues was carried out in a total volume of 250 μL, containing 10 mM Tris-HCl, pH=7.4, 1 mM EDTA and 5 µg MTSEA biotin-XX linker (Biotium) for 30 min at room temperature in the dark. The RNA was then purified by phenolchloroform extraction, denatured by 10 min incubation at 65 °C and further cleaned with 200 μL μMACS Streptavidin MicroBeads suspension (Milentyi). The RNA was incubated with the beads for 15 min at room temperature and the mix was applied to a pre-equilibrated μColumn in the magnetic field of a μMACS magnetic separator. Beads were washed twice with wash buffer (100 mM Tris-HCl, pH=7.4, 10 mM EDTA, 1 M NaCl and 0.1% (v/v) Tween 20). Biotinylated RNA was eluted twice by addition of freshly made 100 mM DTT and cleaned up with RNeasy MinElute kit (QIAGEN), using 1050 μL of 100% ethanol per 200 μL reaction after addition of 700 μL RLT buffer to preserve short RNA fragments.

Libraries for RNA sequencing were prepared using the KAPA RNA HyperPrep Kit (Roche) with minor modifications. 100 ng of RNA per sample were mixed with FPE Buffer, but fragmentation procedure was omitted and RNA was instead denatured at 65 °C for 5 min. The rest of the procedure was performed as recommended by the manufacturer, with the exception of SPRI bead purifications: after adapter ligation, 0.95X and 1X SPRI bead-to-sample volume ratios were used (instead of two rounds of SPRI purification with 0.63X volume ratios) to retain smaller (150-300 bp) cDNA fragments in the library which would otherwise be lost in size selection. The libraries were quality controlled by electrophoresis on a TapeStation System (Agilent), quantified by Qubit (Thermofisher), pooled and sequenced with single end 70 bp reads on a NextSeq2000, with 50,000,000 average reads per sample. Biological triplicates were generated for each condition.

### dxChIP-seq (double-crosslinking chromatin immunoprecipitation and sequencing)

dxChIP-seq was performed essentially as described, with minor modifications^77^. Six 15 cm dishes were seeded per condition, each containing 8.4 × 10^6^ cells. The following day, cells were treated with 20 J/m^2^ dose of UV and allowed to recover for 3 hours. The media was then removed and the cells were quickly washed twice with PBS. After the final wash, 12 mL of 1.66 mM disuccinimidyl glutarate (DSG, ChemCruz) in PBS were added to each plate and incubated at room temperature for 15 min. The DSG solution was then removed, the cells were quickly washed with PBS three times, and 11 mL of freshly prepared PBS solution containing 1% (w/v) formaldehyde, 5 mM HEPES-KOH, pH 7.5, 10 mM NaCl, 0.1 mM EDTA, 50 µM EGTA was added to each plate. After 8 minutes of incubation at room temperature, the reaction was quenched with 1 mL of 1.25 M glycine. After 5 minutes, the cells were washed 3 times with ice-cold PBS, scraped and centrifuged at 2,000 rcf at 4 °C for 7 minutes. The pellet was snap-frozen in liquid nitrogen and stored at –70 °C.

The pellets were quickly thawed at room temperature, resuspended in 30 mL LB1 buffer (50 mM HEPES-KOH, pH 7.5, 140 mM NaCl, 1 mM EDTA, 10% (v/v) Glycerol, 0.5% (v/v) NP-40, 0.25% (v/v) Triton X-100, with addition of protease inhibitors, phosphatase inhibitors and 2 mM NEM) and incubated for 20 minutes while rotating at 4 °C. The cells were then centrifuged at 1,000 rcf at 4 °C for 5 minutes. Each pellet was resuspended in 30 mL of LB2 buffer (10 mM Tris-HCl, pH 8, 200 mM NaCl, 1 mM EDTA, 0.5 mM EGTA, protease inhibitors, phosphatase inhibitors, 2 mM NEM), incubated for 5 minutes at 4 °C and centrifuged at 1,000 rcf at 4 °C for 5 minutes. The pellets containing the chromatin were then resuspended in 4 mL of LB3 (10 mM Tris-HCl, pH 8, 100 mM NaCl, 1 mM EDTA, 0.5 mM EGTA, 0.1% (w/v) freshly added Na-Deoxycholate, 0.5% (w/v) N-lauroylsarcosine, with freshly added phosphatase inhibitors, protease inhibitors and 2 mM NEM) and transferred to 1 mL Covaris tubes (Covaris). Chromatin shearing was performed using a Covaris E220 for 4 minutes with the following settings: PIP = 150, CPB = 1000, Duty factor = 20%, Temperature = 5 °C).

Sheared chromatin was transferred to 2 mL tubes and 200 µL of 10% (v/v) Triton X-100 was added and mixed into each sample before centrifuging at 20,000 rcf at 4 °C for 20 minutes. The supernatant was kept for immunoprecipitation. To balance the chromatin amount among different samples (including treatments and replicates), the double-stranded DNA concentration was measured with Qubit 1X dsDNA high sensitivity kit (Thermo Fisher Scientific) and equal amount of chromatin dsDNA was used for each immunoprecipitation (210 µg in this work). Chromatin was transferred to a new tube, supplemented with 2 µL spike-in chromatin (Active Motif) and precleared with 25 µL protein G dynabeads (Thermo Fisher Scientific), in 0.1% (w/v) BSA solution in PBS, for 1 h at room temperature. Beads were separated from the chromatin using a magnetic separator, and supernatant (pre-cleared chromatin) was transferred to a new protein-LoBind tube (Eppendorf). An aliquot of the pre-cleared chromatin was reserved to use as input and the rest was used for immunoprecipitation. For immunoprecipitation, protein G dynabeads per sample were pre-washed and pre-coated with the desired antibody. Specifically, for Pol II ChIP, 70 µL beads were incubated with 8 μg of D8L4Y antibody (Cell Signaling Technology) and 2 µg spike-in antibody (Active Motif) resuspended in BSA-PBS solution, for 1 h at 4 °C, then washed three times with BSA-PBS solution. For Ser7^P^ ChIP, 70 µL beads were incubated with 15 µg goat antirat IgG and 3 µg spike-in antibody for 1 h at 4 °C, then washed three times with BSA-PBS solution, followed by incubating with 1 mL anti-Ser7^P^ 4E12 serum for 1 h at 4 °C and subsequent washing. Antibody conjugated beads were added to chromatin and the samples were incubated overnight at 4 °C on a turning wheel. The next day, the beads were washed 5 times with ice-cold RIPA buffer (50 mM HEPES-KOH pH=7.5, 500 mM LiCl, 1 mM EDTA, 1% (v/v) NP-40, 0.7% (w/v) freshly added Sodium Deoxycholate) and eluted with 180 µL elution buffer (25 mM Tris-HCl pH=7.5, 5 mM EDTA, 0.5% (w/v) SDS) at 65 °C for 1 hour with shaking. The supernatant was transferred to a new tube and treated with 400 µg Proteinase K (Invitrogen) overnight at 60 °C. The DNA is purified with purification columns (Zymo Research) and libraries were prepared with NEBNext Ultra II DNA Library prep kit (New England Biolabs). The libraries were sequenced with paired end 60 bp reads on a NextSeq2000, with 40,000,000 average reads per sample. Biological triplicates were generated for each condition.

### 3’ End-seq

SWO cells were prepared essentially as described for TT_chem_-seq. RNA was collected using Qiazol (QIAGEN) and purified using the miRNeasy kit (QIAGEN). The purified RNA was quantified using Qubit RNA BR assay (Thermo Fisher Scientific), and 2.5 µg were used with the QuantSeq 3′ mRNA-Seq Library Prep Kit (FWD) (Lexogen, A01172), which uses oligo-dT to prime for first strand synthesis and random priming for second strand synthesis. Samples were spiked with 1% yeast RNA for normalization (see TT_chem_-seq). Libraries were PCR amplified for 12 cycles, quantified by Qubit and quality controlled by electrophoresis on a Bioanalyzer System (Agilent), pooled, and sequenced (2 × 100 bp) on an Illumina NovaSeq 6000.

### GSK3 kinase assay

To assess GSK3 kinase activity on the Pol II CTD, 4 µg of phosphorylated/unphosphorylated peptides were incubated with 4 µg of purified GSK3b^99^, in TBS (20 mM Tris-HCl, pH=7.6, 150 mM NaCl) for 1 hour at room temperature, including 2 µM ATP and 10 mM MgCl_2_. 20 µM of GSK3 inhibitor, CHIR99021, was used as a control to inhibit the kinase activity. Peptide phosphorylation was analyzed by slot blot. Reaction products were applied to a 0.45 µm nitrocellulose membrane through a Bio-Dot SF (slot format) microfiltration unit (Bio-rad).

Blocking and application of antibodies were done as for western blots.

### Nuclear extract preparation

Nuclear extract was obtained using a modified version of the Dignam and Roeder Nuclear Extract protocol^100^. Starting from 2 L of expi-HEK293 cells grown to a density of 3-4 million cells/mL, cells were spun down at 500 rcf, washed with PBS twice and the pellet was resuspended in 5 pellet volumes of Buffer A (10 mM HEPES-KOH, pH 7.9, 10 mM KCl, 1.5 mM MgCl_2_, 0.5 mM polymethanesulfonyl fluoride (PMSF), 0.5 mM DTT) and incubated on ice for 10 min. After incubation the cells were spun at 1,000 rcf for 7 min at 4 °C and the pellet was resuspended in 2 pellet volumes of Buffer A. The suspension was dounced with a loose pestle in a 40 mL homogenizer. Cell membrane rupture was checked with trypan blue under the microscope until greater than 90%. The suspension was centrifuged at 9,000 rcf for 20 min at 4°C. The lipid layer and supernatant were removed. The crude nuclear pellet was resuspended in Buffer C (0.2 mM EDTA, 25% (v/v) glycerol, 20 mM HEPES-KOH, pH 7.9, 1.5 mM MgCl_2_, 0.5 mM PMSF, 0.5 mM DTT) and dounced 10 times with a loose pestle. 3 M KCl was added in a dropwise manner until reaching a final concentration of 0.42 M. The suspension was centrifuged at 35,000 rcf for 1 hour at 4 °C. The supernatant was dialyzed against 40-50 volumes of Buffer D (0.2 mM EDTA, 20% (v/v) glycerol, 20 mM HEPES-KOH, pH 7.9, 0.1 M KCl, 1 mM PMSF, 1 mM DTT) until conductivity was stable, roughly for 5 hours. The dialysate was centrifuged at 12,000 rcf for 20 min at 4 °C and the supernatant was aliquoted, flash-frozen and stored at –70 °C.

### *In vitro* transcription assay

A ∼2 kb long mini gene containing a CMV enhancer, a promoter including TATA box, a GFP coding sequence, and a poly-adenylation and termination signal was immobilized on streptavidin beads through a 5′-biotin functionalized primer and washed twice with Wash Buffer (0.12 mM EDTA, 12% (v/v) glycerol, 12 mM HEPES-KOH, pH7.9, 60mM KCl, 0.0125 % (v/v) NP-40, 7.5 mM MgCl_2_, 0.25 mg/mL BSA, 1 mM DTT). The beads were incubated with nuclear extract for 30 min at 25 °C. After incubation, NTPs were added to initiate transcription, and the reaction was stopped at each timepoint by quickly washing twice with Wash Buffer 2 (0.12 mM EDTA, 12% (v/v) glycerol, 12 mM HEPES-KOH, pH7.9, 60mM KCl, 0.05 % (v/v) NP-40, 9 mM MgCl_2_, 0.6 mM DTT) and boiling in 2x Laemmli Buffer for 5 min at 98 °C. Proteins bound to the DNA were analyzed by western blot. Where applicable, GSK3 and CDK9 inhibitors were added 30 s before adding NTPs.

### Computational Analysis

#### General alignment methods

All reads were trimmed and quality-filtered with Trim Galore, using a quality threshold of 30^101^. Alignment method varied per experiment (see individual sections). Correlation between replicates was checked using deepTools multiBamSummary and bigwigs were created using deepTools bamCoverage^102^.

All reads were aligned to the human genome (hg38) and, where a spike-in was used, to either the *Saccharomyces cerevisiae* (sacCer3) genome or the *Drosophila melanogaster* (dm_r6.46) genome.

Ensembl gene annotation version GRCh38.102 was used to identify human gene type, position and length. Genes were divided into length categories of 0-1000 bp, 1001-5000 bp, 5001-10000 bp, 10001-25000 bp, 25001-50000 bp, 50001-75000 bp and greater than 75000 bp. For 3’Endseq analysis the first three classes were merged into 0-10000 bp, due to the low number of multiple polyA cleavage sites detected in these genes.

For some experiments we further subdivided genes into three classes: 1) protein coding genes – those with an ensembl biotype of protein_coding, 2) snRNA genes – those with an ensembl biotype of snRNA, 3) other ncRNA – those with an ensembl biotype of lncRNA, misc_RNA, miRNA or snoRNA.

Further analysis was performed in R, with visualization using ggplot2^103^.

#### TTchem-seq alignment and processing

Trimmed reads were aligned to the hg38 and sacCer3 genomes using the STAR aligner (v2.6.0a) with basic two-pass mapping^104^. PCR duplicates were marked and removed using Picard^105^. Replicates were merged, and resulting BAM files were split by strand. Bigwig files were normalized to the number of spike-in reads.

#### TTchem-seq metagene profiles

Using ngs.plot (v2.61), genes were split into 100 bins, with size depending on the desired feature^106^. For TSS-centric plots the TSS +/- 5 kb was divided into 100 equal sized bins of 100 bp. For whole-gene plots the gene +/- 5 kb was divided into 100 bins, the 20 bins before the TSS and 20 after the TTS were of equal sizes of 250 bp, while the 60 bins in the gene body were of equal size with length 1/60th of the per gene length. The ngs.plot calculated coverage was normalized to the number of spike-in reads. Confidence intervals were determined as standard error of the mean.

#### Stalling ratio index

To calculate the stalling ratio index we first determined the change in metagene signal between 1/6^th^ of the gene length and 5/6^th^ of the gene length (bins 30 and 70 respectively) for each sample.

Then the change in S7 signal was divided by the change in WT signal.

#### dxChIP-seq alignment and processing

Trimmed reads were aligned to the hg38 and dm_r6.46 genomes using Bowtie2 (v2.5.1) with default parameters^107^, then PCR duplicates were marked and removed with Picard^105^. Replicates were merged and Bigwig files were created normalizing to the total number of reads (Figs. 1 and S1) or number of spike-in reads (Figs. 2 and S2).

We determined the distribution of signal across gene class by counting all reads that aligned within the 500 bp flank from genes of a given class and subtracting this from total signal. This avoids the issue of double counting reads due to overlapping genes within a given class, but not the issue of double counting reads between classes.

#### dxChIP-seq metagene profiles and quantification

Similarly to TTchem-seq metagene profiles, genes were split into 100 bins, with size depending on the desired feature. For TSS-centric plots the TSS +/- 5 kb was divided into 100 equal sized bins of 100 bp each. For whole-gene plots the gene +/- 5 kb was divided into 100 bins, the 20 bins before the TSS and 20 after the TTS were of equal sizes of 250 bp, while the 60 bins in the gene body were of equal size with length 1/60th of the per gene length. Coverage was calculated using bedtools from the already normalized bigwigs and further normalized to average signal 5 kb upstream of all genes. To calculate the ratio of Ser7^P^ signal to total Pol II signal the individual signals were normalized to the input signal, then normalized Ser7^P^ signal was divided by normalized Pol II signal. We plot log2 of the resultant ratio.

For the > 100 kb genes plot in Figure 2I the TSS + 100 kb was divided into 2000 equal size bins of 50 bp each. The relationship of Ser7^P^ signal to total Pol II signal was calculated as above, except that normalized Pol II signal was subtracted from normalized Ser7^P^ signal instead of taking the log2ratio, in order to show data in a similar way as the mathematical modelling predictions in Figure 2H that plotted subtraction.

#### 3’Endseq alignment and processing

Prior to trimming with Trim Galore, 3’ Endseq reads were filtered to contain only those ending with a minimum of 10 As, allowing for one mismatch, using the SeqKit toolkit^108^. These reads were then trimmed and quality-filtered with Trim Galore and the polyA tail removed using cutadapt^101^. Trimmed reads were aligned to the hg38 genome using the STAR aligner with default parameters^104^.

#### Identification of PolyA cleavage sites

PolyA cleavage sites (CS) were identified similarly to Hoffman et al.^109^. Replicates were merged and all coverage of less than 5 reads per bp was discarded. Continuously covered regions were identified as ‘islands’ of coverage and islands within 10 bp were merged. The position of maximum coverage for each island was identified as the peak, where peaks were within 50 bp only the peak with higher coverage was kept. Only peaks with a minimum coverage of 50 reads were used in subsequent analysis. Regions of the genome within 50 bp of these positions were checked for polyA sequences (minimum of 10 consecutive As, allowing for one mismatch) and if one was found the peak was discarded. Finally, we identified peaks contained within genes. The read coverage of these peaks was calculated from the original alignment using samtools bedcov and normalized by total read count. The 84% of peaks within genes were identified as CS.

For genes with at least 2 CS we identified the CS with highest coverage in both WT and S7A samples. We used a two-sample student’s t-test, with Benjamini & Hochberg multiple testing correction and a threshold of 0.05, to determine if these peaks were significantly higher than all other peaks in the gene. If they were we identified if the peak was in the same position in both WT and S7A or not.

### Ser7 hyper-phosphorylation modelling and simulation

We modelled serine 7 hyper-phosphorylation in response to UV damage induced stalling using Beacon Calculus^54^. This system employs process algebra, where each component of the system is treated independently, and the processes are able to interact with other processes simultaneously occurring. Every action is assigned a numerical rate that dictates the speed at which it is performed, with the introduction of inherent randomness as observed in biological systems. We created two models, one for each of two fundamentally different mechanisms of Serine 7 hyperphosphorylation in response to stalling, *in cis* and *in trans* phosphorylation.

Both models were defined with some common components:

Pol II: Individual Pol II molecules were modelled as distinct processes that tracked their position along a linear gene (represented by a parameter i, an integer specifying the location on the gene in units of 100 bp). Each Pol II process also has a parameter p that records its Ser7 phosphorylation state as well as a parameter d that indicates whether the Pol II has met DNA damage.

DNA Damage Generation (MakeDamage[i]): This process introduces damage, modelled by a beacon, at random intervals along the gene which represents UV-induced DNA lesions. The average space between these intervals is given by damage_freq.

DNA Damage Repair (RepairDamage[]): This process removes damage from the gene at a rate determined by the repair_half_life parameter.

Transcription dynamics: The RNA Pol II process was modelled differently pre- and post- transcriptional pausing. Before the pausing site, Pol II moves at a slow rate. Upon reaching the pause_location, Pol II can either terminate transcription and return to the Pol II pool, or enter a transient pause state before proceeding to elongation. Past the pause site, Pol II elongates at a faster rate. When encountering damage at the next base pair, Pol II stalls. Regardless of whether Pol II is stalled or moving, it can dissociate from the gene at a given rate and return to the Pol II pool. Upon reaching the gene end, Pol II unbinds and returns to the pool.

The core distinction between the *cis* and *trans* models lies in the mechanism by which Ser7 is phosphorylated (represented by the internal p flag):

#### *Cis*-Phosphorylation Model

In this model, Ser7 phosphorylation occurs directly on the Pol II that encounters DNA damage. Specifically, within the elongation process, if an unphosphorylated Pol II encounters a damage beacon, its internal p flag is set to 1 at that location and Pol II is stalled until the damage is removed.

#### *Trans*-Phosphorylation Model

In the *trans* model, any Pol II that encounters damage launches a phosphorylation beacon but does not become tagged itself. Critically, for the *trans* mechanism, any Pol II molecule reaching a specific "trans-sensing" location (defined at 200 bp downstream of the TSS) becomes Ser7-phosphorylated (p parameter changes from 0 to 1) if it detects the presence of a phosphorylation beacon anywhere within the gene.

The models are initiated with the MakeDamage[0], a pool of polymerase_count RNAPolII processes, and the RepairDamage[] process running concurrently: MakeDamage[0] || polymerase_count*RNAPolII_pool[] || RepairDamage[];. See Table S1 for a complete list of the parameters used. The simulations were each run 5000 times and profiles of simulated total and Ser7^P^ Pol II occupancy were extracted from the simulation outputs. The source code for the mathematical modelling is available at https://github.com/shutongye/bcs_Pol_II_phosphorylation.

---

**Figure S1.**
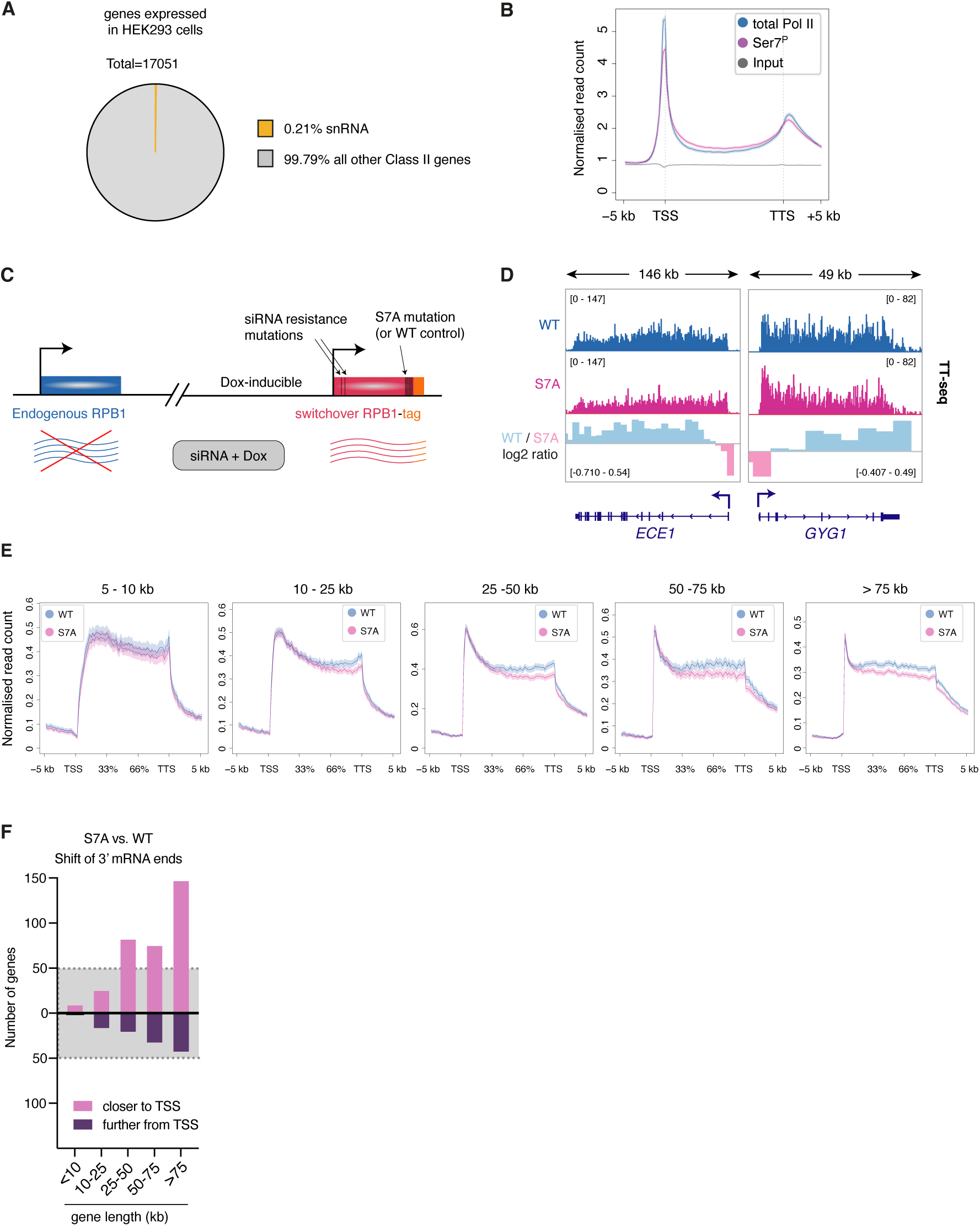
Pol II CTD Ser7 phosphorylation is required for productive elongation. **(A)** Pie chart showing the number of expressed Pol II transcribed (Class II) genes in HEK293 cells (TT_chem_-seq data), separated into two categories: snRNA genes vs. all other genes. **(B)** Metagene plots showing the occupancy of total and Ser7^P^ Pol II on protein-coding genes (relative scale, TSS to TTS), in WT cells. **(C)** Schematic of the switchover system: endogenous RPB1 is silenced using siRNA, while stably integrated, siRNA-resistant and tagged RPB1 is induced with Dox. **(D)** Example of nascent RNA (TT_chem_-seq data) on *ECE1* and *GYG1* genes, in WT and S7A switchover cells. **(E)** Metagene plots of TT_chem_-seq data, showing nascent RNA signal on genes of different length categories, in WT and S7A switchover cells. **(F)** Bar plot showing the number of genes that display different RNA 3’-end usage, in S7A vs. WT switchover cells, separated into different gene length categories.

**Figure S2.**
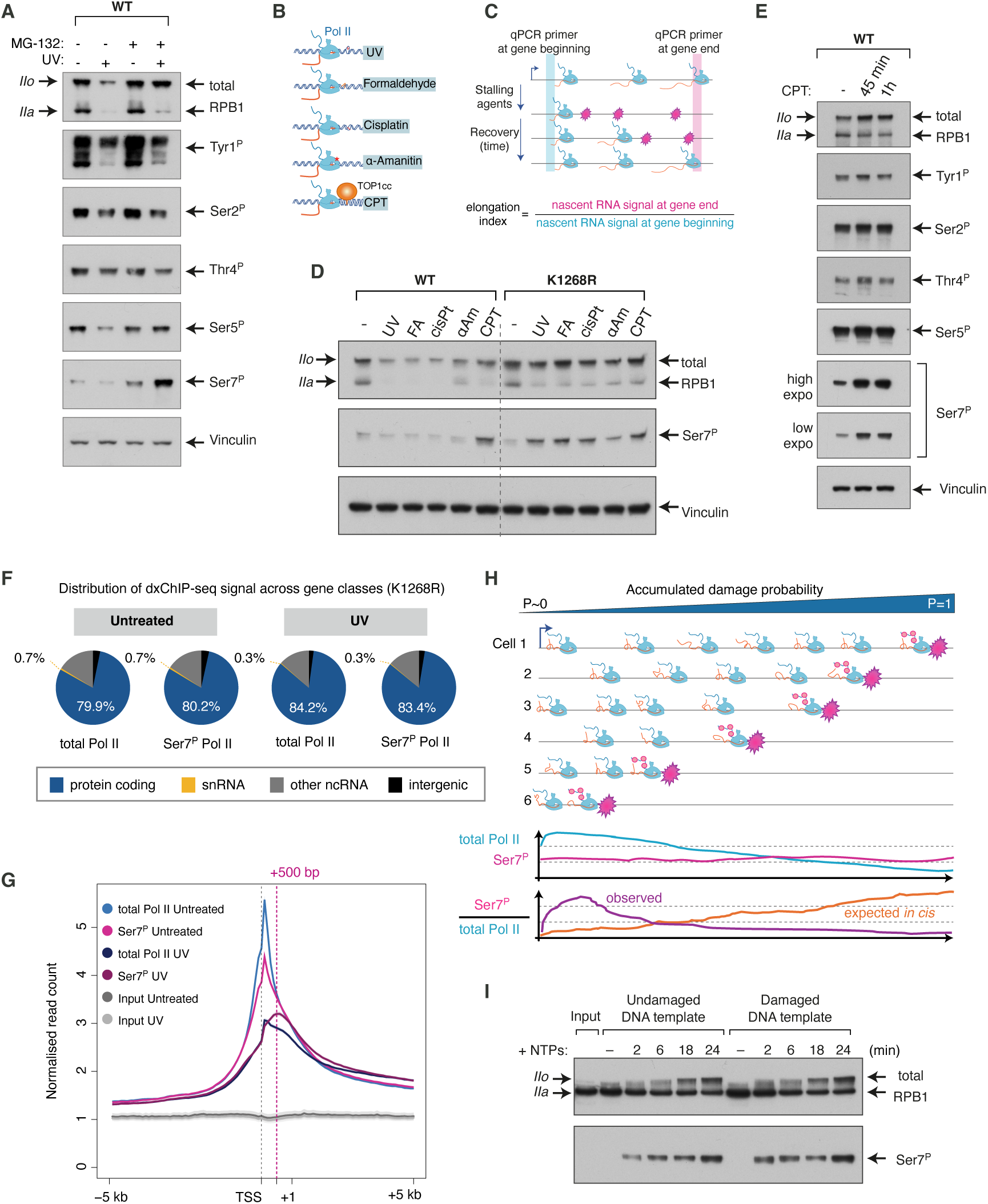
Pol II Ser7 hyper-phosphorylation is a universal hallmark of elongation-stalled transcription, and it occurs *in trans*. **(A)** Western blot detecting the level of total RPB1 and its different phospho-forms in WT HEK293 cells, before and 3 hours post-UV (20 J/m^2^), with or without proteasome inhibitor MG132. **(B)** Sketch illustrating different obstacles on genes caused by a variety of agents. (CPT= Camptothecin, TOP1cc = Topoisomerase 1 cleavage complex). **(C)** Sketch illustrating the principle of the RT-qPCR assay used to derive Pol II elongation index. **(D)** Western blot detecting the total and Ser7^P^ RPB1, in WT and K1268R cells, treated with different agents to induce Pol II stalling. UV = 45 min after UV (20 J/m^2^); FA, formaldehyde = 250 µM for 3 hours; cisPt, cisplatin = 100 µM for 3 hours; αAm, alpha-amanitin = 10 µM for 3 hours; CPT, camptothecin = 20 µM for 3 hours. **(E)** Western blot detecting the level of total RPB1 and its different phospho-forms in WT HEK293 cells, before and after CPT treatment (20 µM). **(F)** Pie charts showing the distribution of total Pol II and Ser7^P^ Pol II signals, detected and quantified by dxChIP-seq, across different gene categories. K1268R cells, before and 3h after UV (20 J/m^2^). **(G)** Metagene plot of dxChIP-seq data, showing the distribution of total and Ser7^P^ Pol II within the first 5 kb from the TSS, in untreated K1268R cells and 3h after UV (20 J/m^2^). **(H)** Schematic illustrating an *in cis* scenario for deposition of Ser7^P^ on stalled Pol II. Below, graphical representations of expected total and Ser7^P^ Pol II occupancy profiles and their ratios. **(I)** *In vitro* transcription assay with nuclear extracts, using either undamaged (left) or UV-damaged (right) DNA template. Total and Ser7^P^ RPB1 were detected using Western blot, at different timepoints after initiating transcription and phosphorylation by adding NTPs.

**Figure S3.**
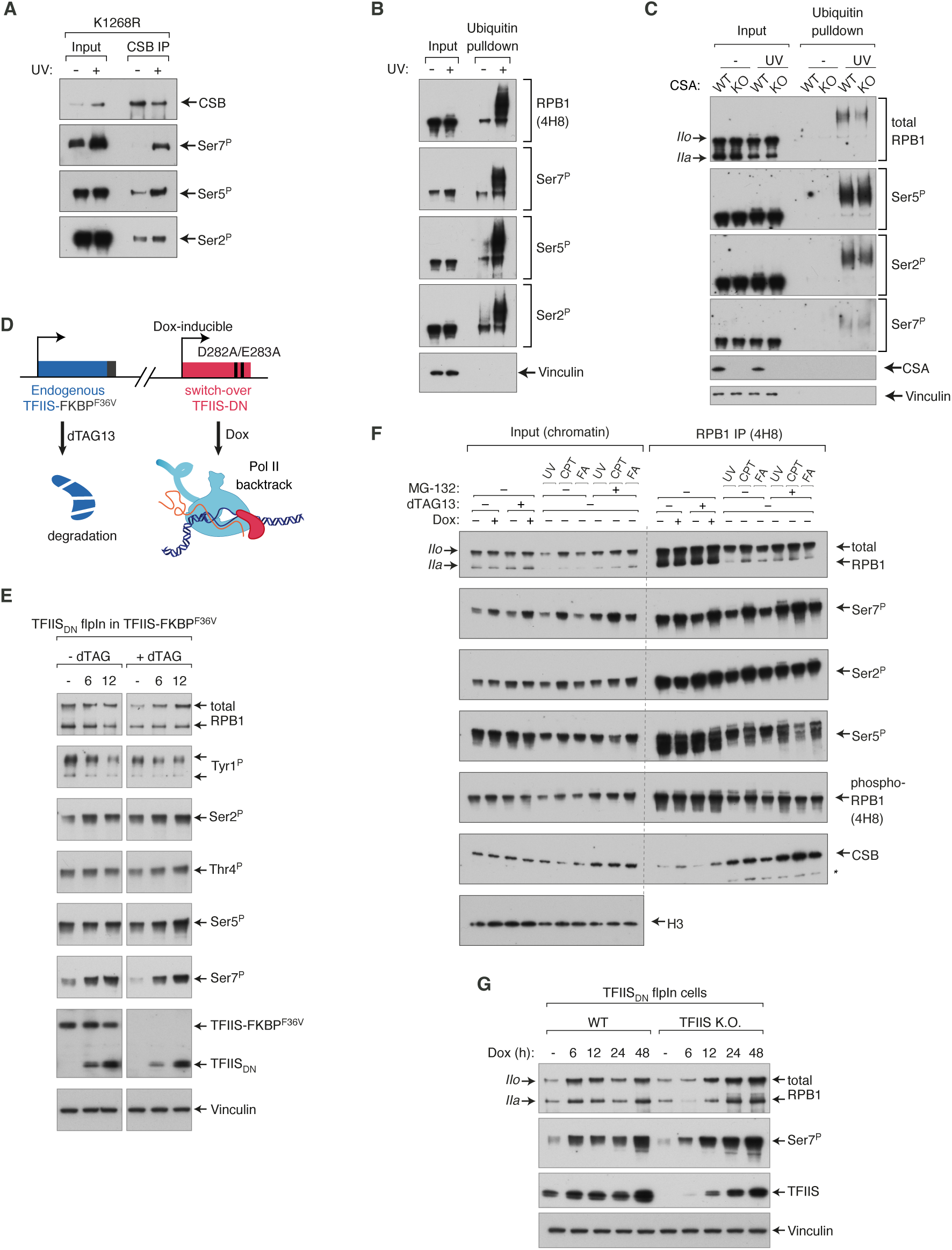
Ser7 hyper-phosphorylation is a separate pathway from TC-NER and the “last resort”, and is triggered by Pol II backtracking. **(A)** Immunoprecipitation of CSB and western blot analysis of different phospho-forms of RPB1, before and 3h after UV treatment (30 J/m^2^), in K1268R cells. **(B)** Ubiquitin pulldown assay followed by western blot detecting total phosphorylated RPB1 and its phospho-forms, before and 45 min post-UV (20 J/m^2^). **(C)** Ubiquitin pulldown assay followed by western blot detecting total RPB1 and its phosphoforms, before and 45 min post-UV (20 J/m^2^), in WT and *CSA* K.O. cells. **(D)** Sketch illustrating TFIIS_DN_ switchover model system. Endogenous, WT TFIIS is tagged with FKBP^F36V^, and upon addition of dTAG13 is rapidly degraded. Simultaneously, expression of TFIIS_DN_ is induced from a stably integrated locus using Dox. **(E)** Western blot detecting total RPB1 and its phospho-forms in TFIIS_DN_ switchover cells, with (left) and without (right) of removal of endogenous WT TFIIS using dTAG. **(F)** Immunoprecipitation of RPB1 (4H8 antibody) from TFIIS_DN_ switchover cells, followed by western blot showing different phospho-forms of RPB1 and CSB, after treatments with stalling agents. Same treatment conditions as Figure 3f. **(G)** Induction of TFIIS_DN_ from a stably integrated locus using Dox, in WT and *TFIIS* K.O. backgrounds. Western blot detecting total RPB1 and Ser7^P^.

**Figure S4.**
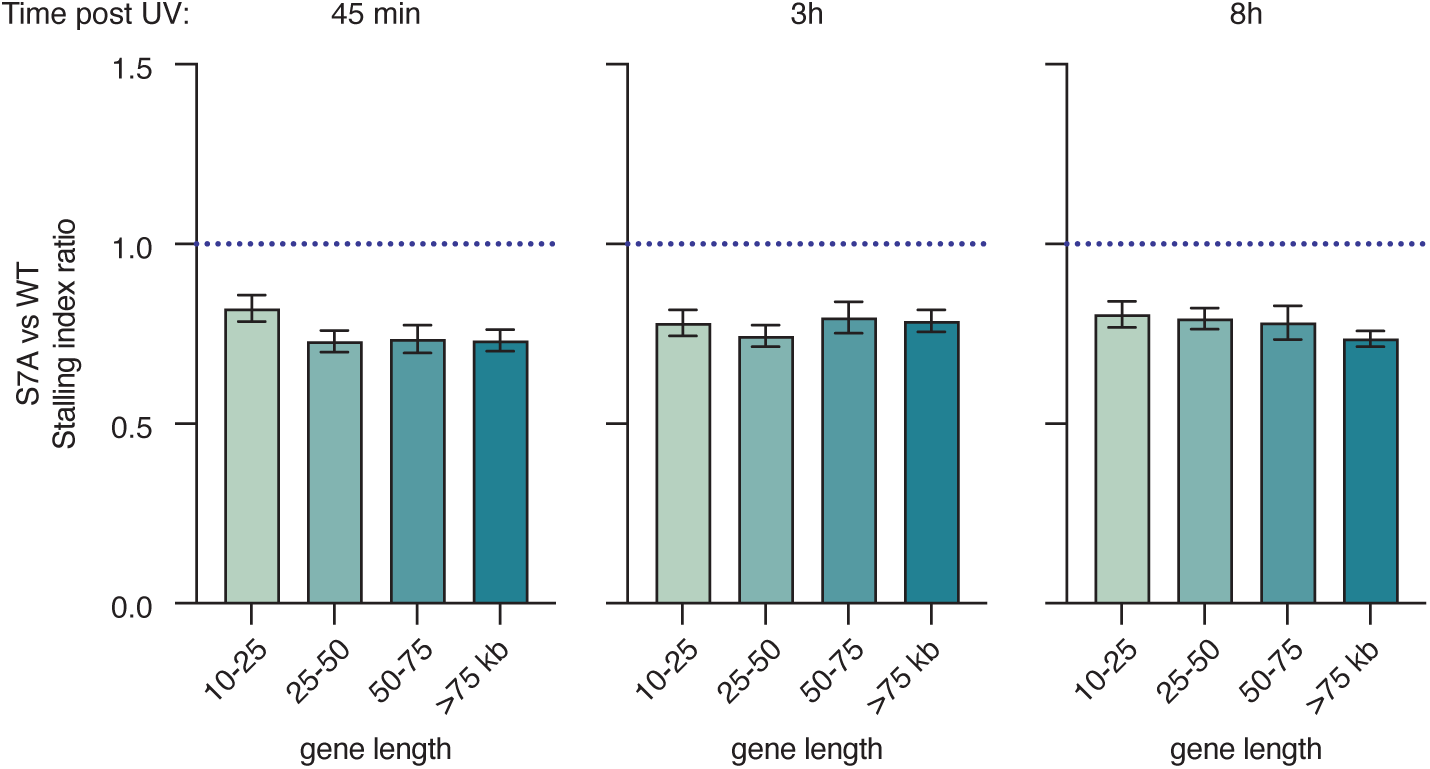
Ser7^P^ is required for transcription recovery after DNA damage. Comparison of stalling indices derived from TT_chem_-seq data, in S7A and WT cells after UV, for different gene length categories. Values above 1 indicate more Pol II stalling in S7A compared to WT cells (untreated condition shown in Figure 1h). All values are statistically significant (p<0.0001, two-tailed t-test).

**Figure S5.**
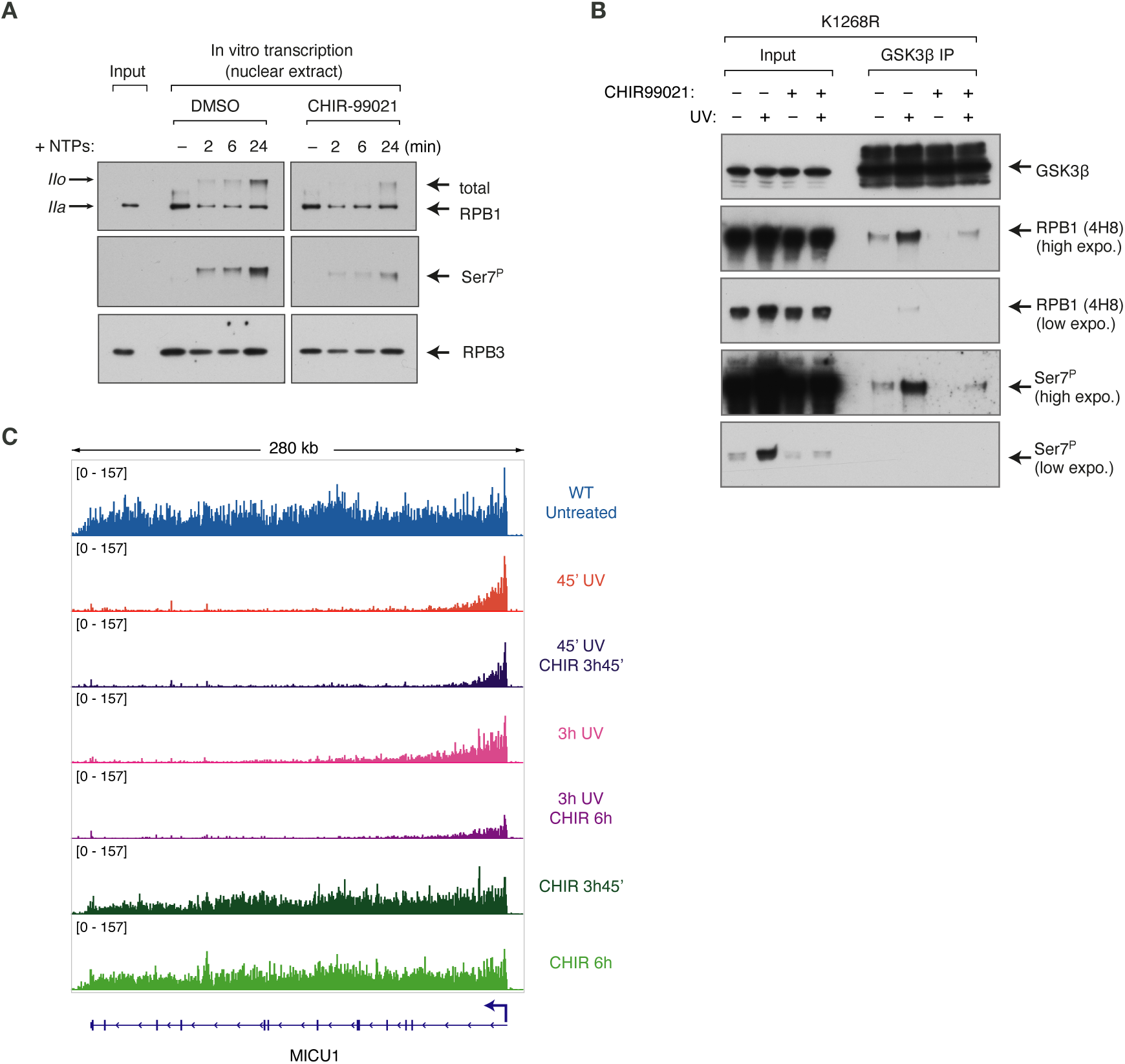
GSK3 is a Pol II Ser7 kinase. **(A)** *In vitro* transcription assay with nuclear extracts followed by western blot detecting total and Ser7^P^ RPB1. Inhibitors of CDK9 (DRB, 100 µM) and GSK3 (CHIR-99021, 10 µM) were added 30 s before initiating transcription and phosphorylation by adding NTPs. RPB3 subunit of Pol II is used as a loading control. **(B)** Immunoprecipitation of GSK3β from chromatin, before and 3h post-UV (20 J/m^2^), in RPB1 K1268R cells. Western blot detects GSK3β, total phosphorylated RPB1 (4H8) and Ser7^P^ RPB1. **(C)** Browser snapshot of an example gene *MICU1*, displaying nascent RNA production (TT_chem_seq data), with or without treatment with CHIR-99021, before and after UV as indicated in the figure.

**Table S1.** Parameters used for the simulation of *in cis* and *in trans* Ser7 phosphorylation.

| Parameter | Value | Unit |
| --- | --- | --- |
| **General** |  |  |
| polymerase_count | 250 | - |
| initiation_freq | 0.4 | 1/s |
| elongation_speed | 33.3 | bp/s |
| **Promoter-Proximal Pausing** |  |  |
| pause_location | 100 | bp |
| pause_dwell_time | 3 | s |
| pause_termination_probability | 0.8 | - |
| pause_release_probability | 0.2 | - |
| **DNA Damage & Repair** |  |  |
| damage_freq | 0.00005 | 1/bp |
| repair_half_life | 14400 | s |
| **Pol II Dissociation & Reuse** |  |  |
| dissociation_time | 600 | s |
| probability_reuse_on_completion | 1 | - |
| **Gene & Simulation Scale** |  |  |
| gene_length | 100000 | bp |
| fast | 1000 | - |

**Table S2.**
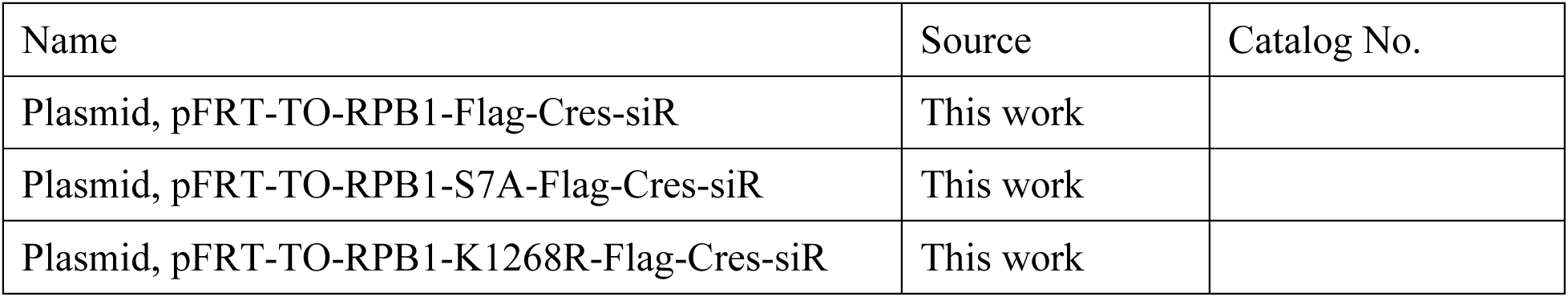
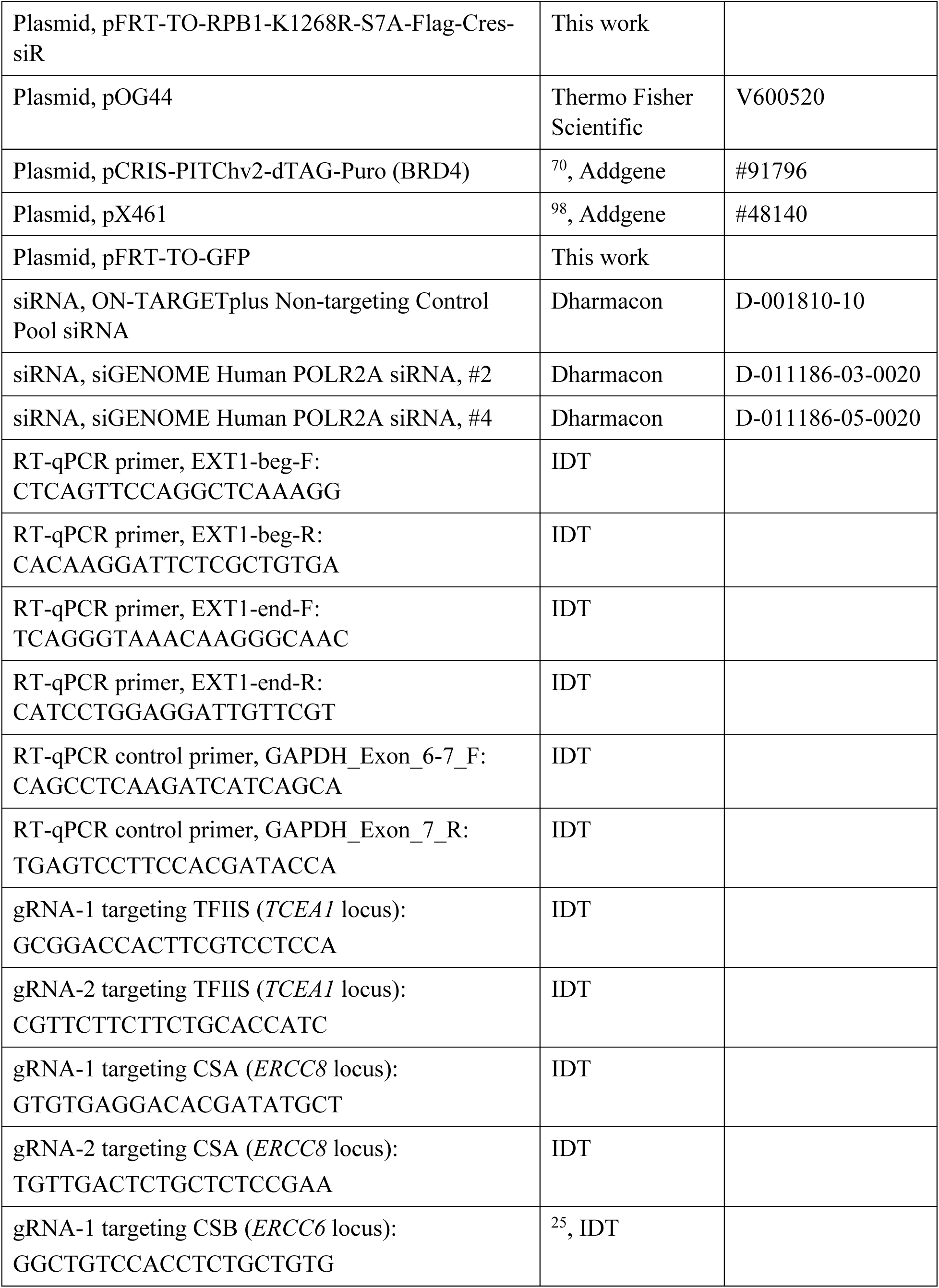
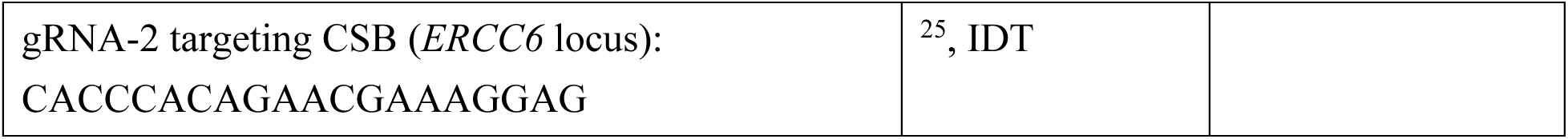
Plasmids and oligonucleotides.

**Table S3.** Cell lines.

| Cell line | Source |
| --- | --- |
| Flp-In T-Rex HEK293 | Thermo Fisher Scientific, R78007 |
| RPB1 K1268R knock-in clone E2 (in Flp-In T-Rex HEK293) | <sup>25</sup> |
| RPB1 K1268R knock-in clone D12 (in Flp-In T-Rex HEK293) | <sup>25</sup> |
| CSB K.O. (in Flp-In T-Rex HEK293) | <sup>25</sup> |
| CSA K.O. (in Flp-In T-Rex HEK293) | This work |
| RPB1 K1268R knock-in + CSB K.O. (in Flp-In T-Rex HEK293) | This work |
| Switchover RPB1-Flag WT (in Flp-In T-Rex HEK293) | This work |
| Switchover RPB1-S7A-Flag (in Flp-In T-Rex HEK293) | This work |
| Switchover RPB1-K1268R-Flag (in Flp-In T-Rex HEK293) | This work |
| Switchover RPB1-K1268R-S7A-Flag (in Flp-In T-Rex HEK293) | This work |
| Switchover TFIIS<sub>DN</sub> (TFIIS-D282A-E283A) in TFIIS-FKBP<sup>F36V</sup> background (in Flp-In T-Rex HEK293) | This work |
| Switchover TFIIS<sub>DN</sub> (TFIIS-D282A-E283A) in TFIIS K.O. background (in Flp-In T-Rex HEK293) | This work |
| Switchover TFIIS<sub>DN</sub> (TFIIS-D282A-E283A) in WT background (in Flp-In T-Rex HEK293) | This work |

**Table S4.**
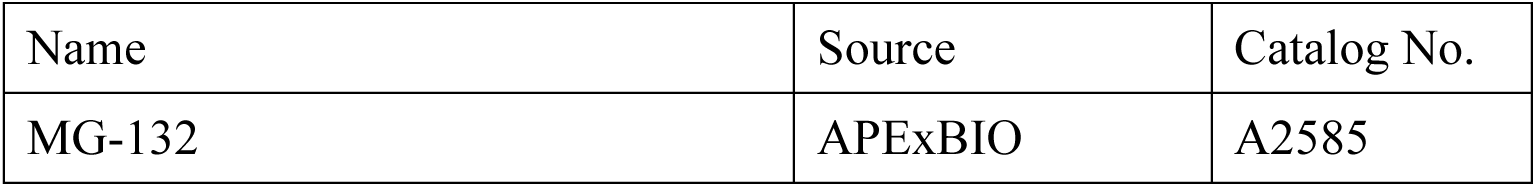
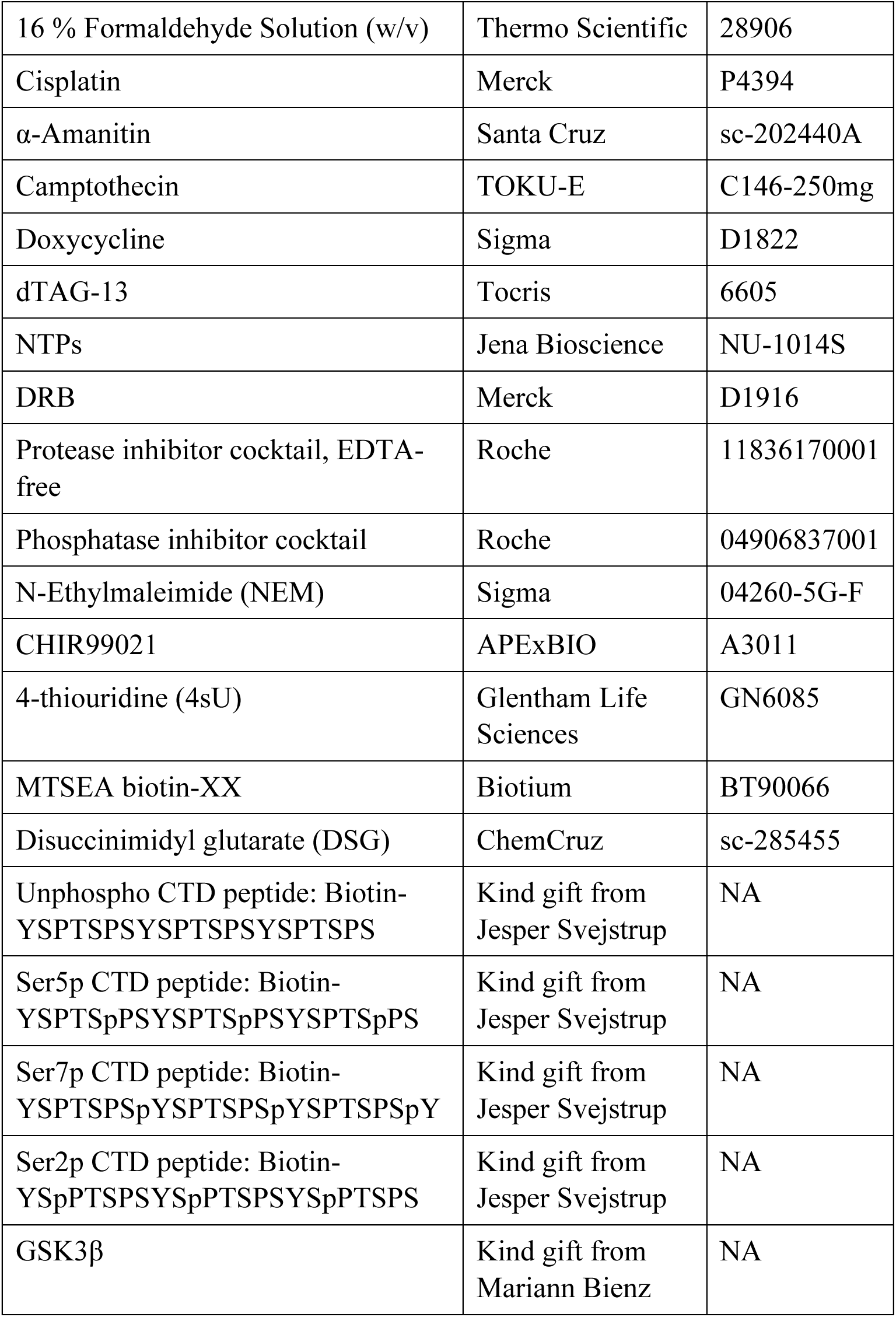
Chemicals, peptides and recombinant proteins.

**Table S5.** Antibodies.

| Antibody | Source | Catalog No. | RRID |
| --- | --- | --- | --- |
| Rabbit monoclonal RPB1 (D8L4Y) | Cell Signaling Technology | 14958S | AB_2687876 |
| Mouse monoclonal RPB1 (4H8) | Abcam | ab5408 | AB_304868 |
| Rat monoclonal RPB1, phospho Ser2 (3E10) | Helmholtz Zentrum Munich | NA | AB_2687450 |
| Rat monoclonal RPB1, phospho Ser5 (3E8) | Helmholtz Zentrum Munich | NA | AB_2687451 |
| Rat monoclonal RPB1, phospho Ser7 (4E12) | Helmholtz Zentrum Munich | NA | AB_2687452 |
| Rat monoclonal RPB1, phospho Tyr1 (3D12) | Helmholtz Zentrum Munich | NA | AB_2793613 |
| Rat monoclonal RPB1, phospho Thr4 (6D7) | Helmholtz Zentrum Munich | NA | AB_2750848 |
| Mouse monoclonal Vinculin | Sigma-Aldrich | V9131 | AB_477629 |
| Rabbit polyclonal CSB/ERCC6 | Bethyl Lab | A301-345A | AB_937849 |
| Rabbit polyclonal CSA/ERCC8 | Abcam | ab137033 | AB_2783825 |
| Rabbit polyclonal TFIIS | Bethyl Lab | A302-240A | AB_1730887 |
| Rabbit polyclonal DYKDDDDK Tag antibody | Cell Signaling Technology | 2368S | AB_2217020 |
| Rabbit polyclonal histone H3 antibody | Cell Signaling Technology | 9715S | AB_331563 |
| Rabbit polyclonal anti-gamma H2A.X (phospho S139) | Abcam | ab2893 | AB_303388 |
| Rabbit polyclonal GSK-3β antibody | Cell Signaling Technology | 9315S | AB_490890 |
| Rabbit polyclonal RPB3 | Bethyl Lab | A303-771A | AB_11218388 |
| Peroxidase-AffiniPure Goat Anti-Rat IgG (H+L) | Jackson ImmunoResearch | 112-035-143 | AB_2338138 |
| Polyclonal Swine Anti-Rabbit Immunoglobulins/HRP | Agilent | P0399 | AB_2617141 |
| Polyclonal Goat Anti-Mouse Immunoglobulins/HRP | Agilent | P0447 | AB_2617137 |

